# Synergistic stabilization of microtubules by BUB-1, HCP-1 and CLS-2 controls meiotic spindle assembly in *C. elegans* oocytes

**DOI:** 10.1101/2022.08.24.505151

**Authors:** Nicolas Macaisne, Laura Bellutti, Kimberley Laband, Frances Edwards, Laras Pitayu-Nugroho, Alison Gervais, Thadshagine Ganeswaran, Hélène Geoffroy, Gilliane Maton, Julie C. Canman, Benjamin Lacroix, Julien Dumont

**Author notes:** Equal contribution (Nicolas Macaisne, Laura Bellutti, Kimberley Laband, Frances Edwards). Corresponding authors: Julien Dumont, 15 rue Hélène Brion-254B, 75205 Paris Cedex 13, France; Benjamin Lacroix, 1919, route de Mende, 34293 Montpellier Cedex 5, France.

## Abstract

During cell division, chromosome segregation is orchestrated by a microtubule-based spindle. Interaction between spindle microtubules and kinetochores is central to the bi-orientation of chromosomes. Initially dynamic to allow spindle assembly and kinetochore attachments, which is essential for chromosome alignment, microtubules are eventually stabilized for efficient segregation of sister chromatids and homologous chromosomes during mitosis and meiosis I respectively. Therefore, the precise control of microtubule dynamics is of utmost importance during mitosis and meiosis. Here, we study the assembly and role of a kinetochore module, comprised of the kinase BUB-1, the two redundant CENP-F orthologs HCP-1/2, and the CLASP family member CLS-2 (hereafter termed the BHC module), in the control of microtubule dynamics in *Caenorhabditis elegans* oocytes. Using a combination of *in vivo* structure-function analyses of BHC components and *in vitro* microtubule-based assays, we show that BHC components stabilize microtubules, which is essential for meiotic spindle formation and accurate chromosome segregation. Overall, our results show that BUB-1 and HCP-1/2 do not only act as targeting components for CLS-2 at kinetochores, but also synergistically control kinetochore-microtubule dynamics by promoting microtubule pause. Together, our results suggest that BUB-1 and HCP-1/2 actively participate in the control of kinetochore-microtubule dynamics in the context of the BHC module to promote accurate chromosome segregation in meiosis.

## INTRODUCTION

Equal partitioning of the replicated genome between the two daughter cells is a key step of cell division. Throughout meiosis and mitosis, proper interactions between spindle microtubules and kinetochores, multiprotein complexes assembled at the centromeres of meiotic and mitotic chromosomes, are essential for accurate chromosome segregation ^1^. Kinetochore-microtubule attachments are required for co-orientation of sister chromosomes attached to the same spindle pole during meiosis I and for chromosome bi-orientation with sister chromatids attached to microtubules emanating from opposite spindle poles in meiosis II and mitosis ^2^.

After nuclear envelop breakdown (NEBD), dynamic microtubules grow toward the chromosomes where they engage in lateral interactions with kinetochore-localized motor proteins ^3–8^. These initial interactions promote chromosome orientation and accelerate stable end-on attachments with kinetochore-microtubules mediated by the Ndc80 complex ^9–13^. Initially spindle microtubules are highly dynamic to allow spindle assembly and capture by kinetochores, which is essential for chromosome alignment. But as meiosis and mitosis progress, kinetochore microtubules become stabilized for efficient segregation of sister chromatids and homologous chromosomes ^14–16^. Thus, precise control of microtubule dynamics is essential for spindle assembly and the stepwise attachment of chromosomes followed by their accurate segregation.

Proteins of the cytoplasmic linker-associated protein (CLASP) family are evolutionary-conserved regulators of microtubule dynamics ^17–19^. During meiosis and mitosis, CLASP proteins prevent spindle abnormalities and chromosome segregation errors in most species including yeast, *Drosophila*, *C. elegans* and mammals ^20–23^. *In vitro*, CLASPs maintain microtubules in a growing state by promoting microtubule rescue while inhibiting catastrophe ^24–28^. In dividing human cells, two paralogous CLASP1/2 proteins act redundantly at the kinetochore where they are targeted through their C-terminal domain (CTD) by a poorly characterized pathway that involves the motor protein CENP-E and the kinetochore and spindle-associated protein SPAG5/Astrin ^29–33^. In *C. elegans*, CLS-2 is the sole CLASP ortholog that localizes at the kinetochore and is essential for normal spindle assembly and chromosome segregation ^22^. During meiosis in *C. elegans* oocytes, CLS-2 is essential for meiotic spindle assembly, chromosome segregation and polar body extrusion ^34–37^. Kinetochore localization of CLS-2 requires interaction with the two CENP-F-like proteins HCP-1/2, which are themselves localized downstream of BUB-1 (Figure 1A) ^22, 38, 39^.

**Figure 1.**
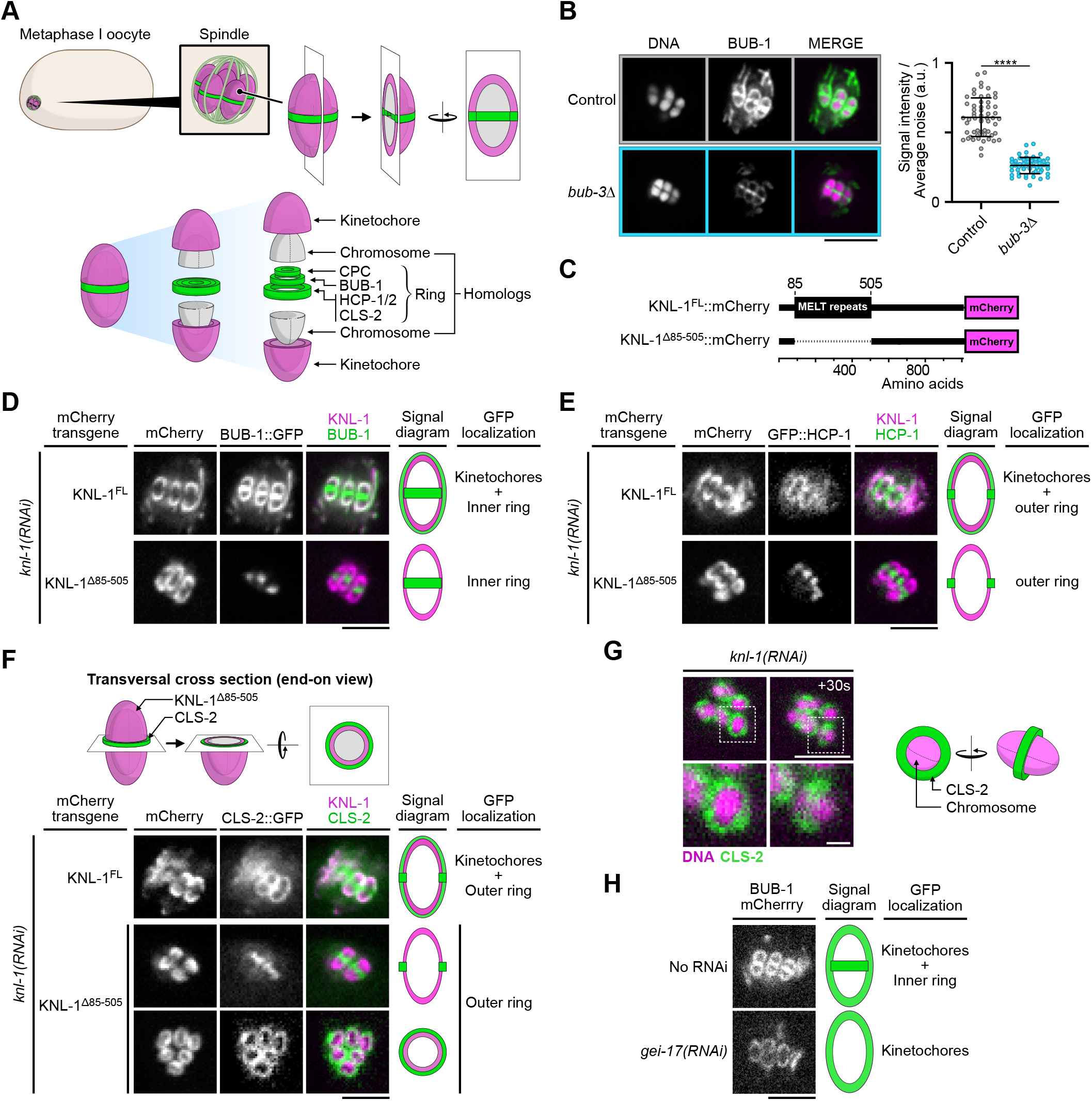
The N-terminal MELT repeats of KNL-1, but not BUB-3, are required for BHC module kinetochore targeting in oocytes. (A) Schematics of kinetochore and ring domain protein organization around a bivalent chromosome during metaphase I in a *C. elegans* oocyte. (B) Immunolocalization of BUB-1 (left) and quantification of BUB-1 signal at kinetochores (right) in *bub-3(ok3437)* mutants *(bub3Δ,* n=58) compared to wild type controls (n=58). Unpaired t-test, aplha=0.05, P<0.0001. (C) Schematics of KNL-1::mCherry protein fusions. (D-F) Localization of BUB-1::GFP (D), GFP::HCP-1 (E) and CLS-2::GFP (F) in worms carrying full length or MELT-deleted KNL-1::mCherry (KNL-1^FL^ and KNL-1^Δ 85-505^ respectively, n≥10). (G) Localization of CLS-2::GFP at ring domains in *knl-1*-depleted oocytes (left) with corresponding schematics (right). (H) Localization of BUB-1::mCherry in *gei-17*-depleted oocytes (n=29) compared to controls (n=25). Scale bars 5 µm, 1 µm in insets.

Bub1 (BUB-1 in *C. elegans*) is a kinase originally identified for its role in the Spindle Assembly Checkpoint (SAC), a safety mechanism that ensures proper connection of kinetochores to spindle microtubules ^40, 41^. During meiosis and mitosis, Bub1 is also directly involved in chromosome bi-orientation through its non-SAC functions that: 1) promote kinetochore recruitment of dynein and CENP-F, 2) ensure proper inner-centromere localization of Aurora B, 3) recruit PP2A:B56 on meiotic chromosomes, and 4) limit kinetochore-microtubule attachment maturation by the SKA complex in mitosis ^38, 39, 42–48^. During mitosis, Bub1 interacts physically with Bub3 via its ‘Bub3-binding motif’, formerly known as the GLEBS domain ^49–51^. The Bub1/Bub3 complex is then recruited to kinetochores through Bub3 direct-binding to phosphorylated MELT (Met-Glu-Leu-Thr) repeats located in the N-terminal half of Knl1 ^51–56^. During meiosis in *C. elegans* oocytes, BUB-1 also localizes to kinetochores, which display characteristic cup-like shapes ^57^. This kinetochore localization requires KNL-1, but whether it occurs via BUB-3 and the KNL-1 MELT repeats is unknown ^34^. BUB-1 additionally concentrates at trilaminar ring domains located between each pair of homologous chromosomes in meiosis I and between sisters in meiosis II ^34^. This localization lies downstream of the Chromosomal Passenger Complex (CPC) and requires BUB-1 sumoylation ^36, 58, 59^. The specific functions of these two distinct chromosomal BUB-1 localizations are at present unclear. However, BUB-1 plays a critical role during oocyte meiosis as its depletion leads to severe chromosome segregation errors and meiotic spindle abnormalities ^34^. These meiotic phenotypes have been attributed to the role of BUB-1 in the recruitment of HCP-1/2 and CLS-2 at kinetochores and ring domains, the SUMO-dependent targeting of the chromokinesin KLP-19, and the phospho-dependent recruitment of PP2A:B56 to ring domains ^34, 48, 59, 60^. However, the exact function of BUB-1 during meiosis in oocytes is unclear.

In mammals, CENP-F is a large coiled-coil protein recruited to kinetochores through the physical interaction between a specific targeting domain and the kinase domain of Bub1 ^46, 61^. CENP-F contains two high-affinity microtubule binding domains (MTBDs), located at either terminus of the protein, required for the generation of normal interkinetochore tension and stable kinetochore-microtubule attachments ^62–65^. CENP-F is also involved in the recruitment of the dynein motor at kinetochores via its direct interaction with the NudE/L dynein adaptor proteins ^66–70^. Despite these contributions to the process of chromosome alignment, CENP-F is non-essential in mammals as evidenced by the lack of segregation defects in CRISPR knockouts in human cells and by the viability of CENP-F knockout mice ^71–73^. In contrast in *C. elegans*, the two CENP-F-like proteins HCP-1/2 are essential for embryonic viability ^22^. Together with their downstream partner CLS-2, kinetochore-localized HCP-1/2 control kinetochore-microtubule dynamics to prevent sister chromatid co-segregation to the same spindle pole in mitosis, and promote midzone microtubule assembly for central spindle formation in anaphase ^22, 74–76^. During meiosis in *C. elegans* oocytes, HCP-1/2 and CLS-2 localize along spindle microtubules, at the cup-shaped kinetochores and on the ring domains ^34, 35^. Together they are essential for the formation of bipolar meiotic acentrosomal spindles in oocytes, for meiotic chromosome segregation and for efficient cytokinesis during polar body extrusion ^34–37^. Therefore, BUB-1, HCP-1/2 and CLS-2, hereafter termed the BHC module, form a kinetochore module essential for meiosis and mitosis. How is this kinetochore module assembled and whether BUB-1 and HCP-1/2 only act as CLS-2 kinetochore-targeting subunits or whether they participate in regulating kinetochore microtubule dynamics is unknown.

In this study, we investigated the assembly and function of the BHC kinetochore module in the *C. elegans* oocyte and 1-cell embryo. Through a combination of genetic approaches and live cell imaging, we found that the BUB-1 kinase domain, specific C- and N-terminal sequences of HCP-1, and the CLS-2 CTD are essential for BHC module assembly. Our *in vitro* TIRF (Total Internal Reflection Microscopy)-based microtubule assays demonstrated that BHC module components decreased the catastrophe frequency, while increasing the growth rate and rescue frequency. Surprisingly, they also induced a strong synergistic increase of the time spent in pause by microtubules. Overall, our results suggest that BUB-1 and HCP-1/2 are not mere targeting components for CLS-2 at the kinetochore, but instead actively participate in the control of kinetochore-microtubule dynamics in the context of the BHC module, which is essential for accurate chromosome segregation.

## RESULTS

### The N-terminal MELT repeats of KNL-1, but not BUB-3, are essential for BHC module kinetochore targeting in oocytes (Figure 1)

We first determined the molecular mechanisms that target the BHC module to kinetochores and ring domains in *C. elegans* oocytes. In most systems during mitosis, Bub1 localization to kinetochores requires a physical interaction with Bub3, which in turn interacts with the phosphorylated MELT repeats of KNL-1 ^51–56, 77^. We first analyzed if this was the case in *C. elegans* by analyzing a viable *bub-3* deletion mutant (*bub-3(ok3437)*, referred to as *bub-3Δ*). As previously shown, BUB-1 is destabilized in this mutant, with its overall protein levels down to below 10% compared to control worms ^78^. Accordingly, BUB-1 was also significantly reduced at meiotic kinetochores in this mutant, albeit to a much lower extent (55% reduction on average compared to controls) than the overall protein level reduction (Figure 1B). This surprisingly suggested that, in contrast to in yeasts or mammalian cells, BUB-1 could be recruited at kinetochores independently of BUB-3 in *C. elegans* oocytes.

We next tested if the KNL-1 MELT repeats are required for BUB-1 kinetochore targeting in meiosis by using a *C. elegans* strain expressing a truncation mutant of KNL-1 (KNL-1^Δ85-505^) that lacks all MELT repeats ^78, 79^ (Figure 1C). We analyzed the localization of GFP-tagged BUB-1 in strains expressing RNAi-resistant transgenes encoding wild-type KNL-1 (KNL-1^WT^) or KNL-1^Δ85-505^ after depletion of endogenous KNL-1 by RNAi treatment. As expected, in the presence of KNL-1^WT^, BUB-1 localized to cup-shaped kinetochores and to ring domains. In contrast in the presence of KNL-1^Δ85-505^, BUB-1 failed to localize to kinetochores (Figure 1D). GFP-tagged HCP-1 and CLS-2 also did not localize to kinetochores in KNL-1^Δ85-505^ (Figure 1 E, F). The remaining chromosomal signal of GFP-tagged HCP-1 and CLS-2 in this mutant corresponded to their KNL-1-independent ring domain localization, as evidenced by the identical GFP pattern observed in KNL-1-depleted oocytes (Figure 1G). Thus kinetochore, but not ring-domain, localization of BUB-1 depends on the KNL-1 MELT repeats. In line with a previous study, we also confirmed that BUB-1 ring domain localization required the E3 SUMO-protein ligase GEI-17, and thus probably BUB-1 sumoylation (Figure 1H) ^58^. Overall, these results show that in *C. elegans* oocytes, the BHC module can be recruited independently of BUB-3 to kinetochores, but not to ring domains, via the KNL-1 MELT repeats.

### Molecular determinants of BHC module assembly (Figure 2)

Next, we analyzed the domains of BUB-1, HCP-1, and CLS-2 necessary for BHC module assembly. During mitosis, CENP-F in mammals and HCP-1 in *C. elegans*, are recruited to kinetochores through the BUB-1 kinase domain ^39, 46^. We first tested if that was also the case during meiosis by analyzing HCP-1 and CLS-2 localization in *C. elegans* oocytes expressing RNAi-resistant transgenes encoding full length BUB-1 (BUB-1^FL^) or a kinase domain-deleted mutant of BUB-1 (BUB-1^ΔKD^) after depletion of endogenous BUB-1 (Figure 2A) ^39^. GFP-tagged HCP-1 (and CLS-2) localized to kinetochores and ring domains in the presence of BUB-1^FL^, but not BUB-1^ΔKD^ (Figure 2B, C). To determine if the kinase activity of BUB-1 was required for HCP-1 and CLS-2 kinetochore and ring targeting, we also analyzed their localizations in oocytes expressing a kinase dead (BUB-1^D814N^) version of BUB-1 ^79^. Upon depletion of endogenous BUB-1 in this mutant, HCP-1 (and CLS-2) were normally targeted to kinetochores and rings (Figure 2B, C). Therefore, BUB-1 recruits HCP-1 (and CLS-2) to kinetochores and rings through its kinase domain, independently of kinase activity, in meiosis in *C. elegans*.

**Figure 2.**
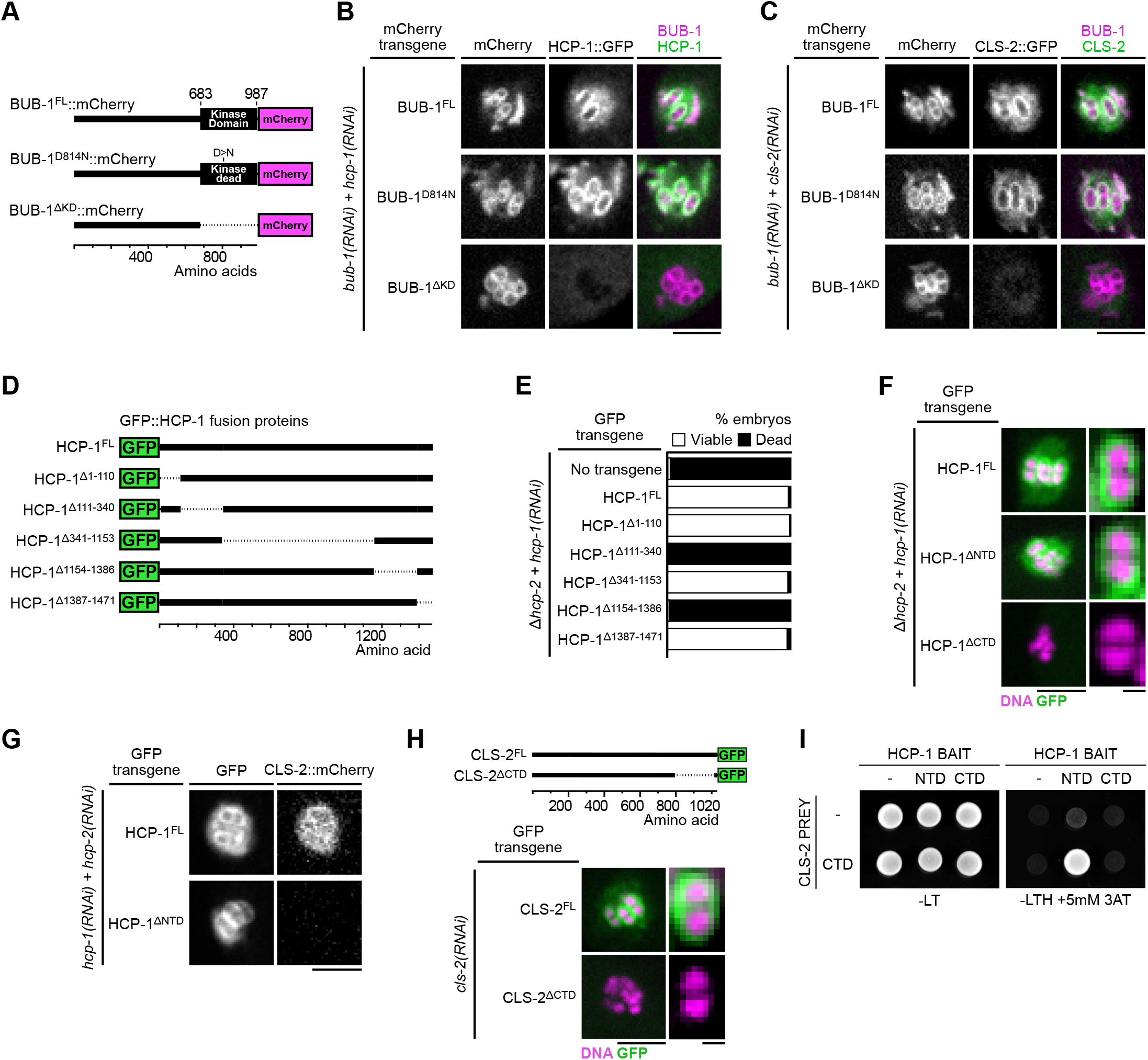
Molecular determinants of BHC module assembly. (A) Schematics of BUB-1::mCherry protein fusion. (B,C) Localization of GFP::HCP-1 (B) and CLS-2::GFP (C) in indicated conditions (n≥10). (D) Schematics of truncated GFP::HCP-1 fusions. (E) Embryonic viability assay in indicated transgenic *hcp-2(ĳm6) (Δhcp-2)* mutants upon depletion of endogenous *hcp-1*. (F) Localization of indicated GFP::HCP-1 fusions in *Δhcp-2* worms depleted of endogenous *hcp-1* (n≥9). (G) Localization of CLS-2::mCherry in indicated conditions (n≥13). (H) Localization of schematized (top) RNAi-resistant CLS-2::GFP fusions upon depletion of endogenous *cls-2* (bottom, n≥10). (I) Yeast two hybrid interaction assay between HCP-1 domains (baits) and CLS-2 CTD (prey). Scale bars, metaphase plate 5 µm, single chromosome details 1 µm.

In mammals, CENP-F kinetochore targeting involves physical interaction between a specific C-terminal CENP-F targeting domain and the kinase domain of Bub1 ^46^. HCP-1 and 2 are long coiled-coil proteins that only share 27.5% and 24.7% similarity respectively with CENP-F, which precluded identification of conserved functional domains ^76^. To identify the domains of HCP-1 responsible for its localization, we thus generated a series of transgenic strains expressing RNAi-resistant GFP-fused truncations of HCP-1 (Figure 2D, figure 2 – Figure supplement 1A). We decided to specifically focus on HCP-1 because, although HCP-1 and 2 are functionally redundant and can compensate for each other to support embryonic development, HCP-1 plays a more primary role in early embryos ^76^. We introduced the HCP-1 transgenes in an *hcp-2* deletion mutant (hereafter *hcp-2Δ*) and analyzed embryonic viability after endogenous HCP-1 depletion by RNAi ^39^. We first verified that GFP-fused wild-type HCP-1 could rescue embryonic lethality in absence of endogenous HCP-1/2. Two main regions of HCP-1 (amino acid 111-340 and 1154-1386, hereafter referred to as NTD and CTD respectively), one at either terminus, were essential for embryonic viability (Figure 2E, figure 2 – Figure supplement 1B), even though the mutant and wild-type transgenes were expressed at comparable levels (Figure 2 – Figure supplement 1C). Although this domain conformation is reminiscent of the 2 MTBDs of human CENP-F, we could not identify significant sequence or structural similarity between the human and *C. elegans* domains ^64^.

We next tested if these C- and N-terminal domains could instead be important for kinetochore localization of the corresponding GFP-fused truncated proteins. We found that deleting the CTD (HCP-1^ΔCTD^), but not the NTD (HCP-1^ΔNTD^), prevented kinetochore targeting of the corresponding deletion (Figure 2F). The CTD could thus be responsible for kinetochore targeting via binding to the kinase domain of BUB-1, although we were unable to confirm direct interaction with a yeast-two-hybrid assay between the HCP-1 CTD and full-length BUB-1 or the BUB-1 kinase domain (Figure 2 – Figure supplement 1D). If the NTD is not required for HCP-1 kinetochore targeting, embryonic lethality in the corresponding mutant, could instead be caused by a defect in CLS-2 recruitment to kinetochores. Accordingly, we found that the HCP-1 NTD truncation mutant prevented mCherry-fused CLS-2 kinetochore targeting (Figure 2G, Figure 2 - Figure supplement 1E). We also found identical results for HCP-1 and CLS-2 localizations in mitosis (Figure 2 – Figure supplement 1F). Thus HCP-1 kinetochore targeting requires its CTD, while CLS-2 kinetochore recruitment is mediated by the NTD of HCP-1.

We then focused on the CLS-2 domain that interacts with the HCP-1 NTD and essential for its kinetochore targeting. In most species, CLASPs targeting to their various subcellular localizations, including to kinetochores, relies on interactions with various adapter proteins via a C-terminal domain (CTD) ^29^. To test the function of the CLS-2 CTD, we generated a transgenic strain expressing a GFP-tagged deletion of the CTD (CLS-2^ΔCTD^). We compared the localization of the corresponding protein to that of full length GFP-fused CLS-2 (CLS-2^FL^) following depletion of endogenous CLS-2. Both transgenes were expressed at similar levels. In contrast to CLS-2^FL^, the CTD-deleted transgenic protein did not target to the kinetochores during meiosis in oocytes or in mitosis in zygotes (Figure 2H, Figure 5 – Figure supplement 1E), which is in line with previous findings on vertebrate CLASPs ^29^. We next performed a yeast two-hybrid-based assay between the CLS-2 CTD and the HCP-1 NTD, which confirmed that the two domains directly interacted to promote CLS-2 kinetochore targeting (Figure 2I, Figure 2 – Figure supplement 1G). Overall, our results show that the BHC module assembly in meiosis and mitosis requires the BUB-1 kinase domain and the HCP-1 CTD, and involves HCP-1 NTD binding to the CLS-2 CTD.

### Kinetochore and ring domain pools of the BHC module act redundantly in spindle assembly and chromosome segregation in oocytes (Figure 3)

We next tested the effect of delocalizing the BHC module from kinetochores in meiosis by preventing its recruitment in KNL-1^Δ85-505^ mutant oocytes. For this we performed live imaging to monitor spindle assembly and chromosome segregation in a strain expressing H2B::mCherry and GFP::β-Tubulin to label chromosomes and microtubules respectively. Although depletion of endogenous KNL-1 in KNL-1^Δ85-505^ mutant oocytes did not prevent assembly of bipolar spindles, but the spindles were smaller and displayed a reduced microtubule density (Figure 3A, B), demonstrating that BHC module kinetochore targeting is essential for normal spindle assembly. These shorter KNL-1^Δ85-505^ mutant spindles were however capable of efficient chromosome segregation, unlike spindles assembled in the complete absence of KNL-1 (Figure 3A-B, Movie 1, Movie 2, Figure 3 - Figure supplement 1). The KNL-1^Δ85-505^ mutant spindles also contrasted with oocytes fully depleted of BUB-1, which displayed severe spindle abnormalities and chromosome segregation defects (Figure 3A, Movie 1, Movie 2, Figure 3 – Figure supplement 1). In control oocytes with endogenous wild-type KNL-1, BHC module components localized at kinetochores and at ring domains. In KNL-1^Δ85-505^ mutant oocytes, the BHC module still targeted to ring domains downstream of GEI-17-dependent BUB-1 sumoylation (Figure 1D-H). Yet in the presence of endogenous wildtype KNL-1, GEI-17-depleted oocytes formed normal bipolar spindles and chromosome segregation was accurate (Figure 3A, Movie 2). We thus hypothesized that the kinetochore and ring domain pools of BHC module could act redundantly for spindle assembly and chromosome segregation in oocytes.

**Figure 3.**
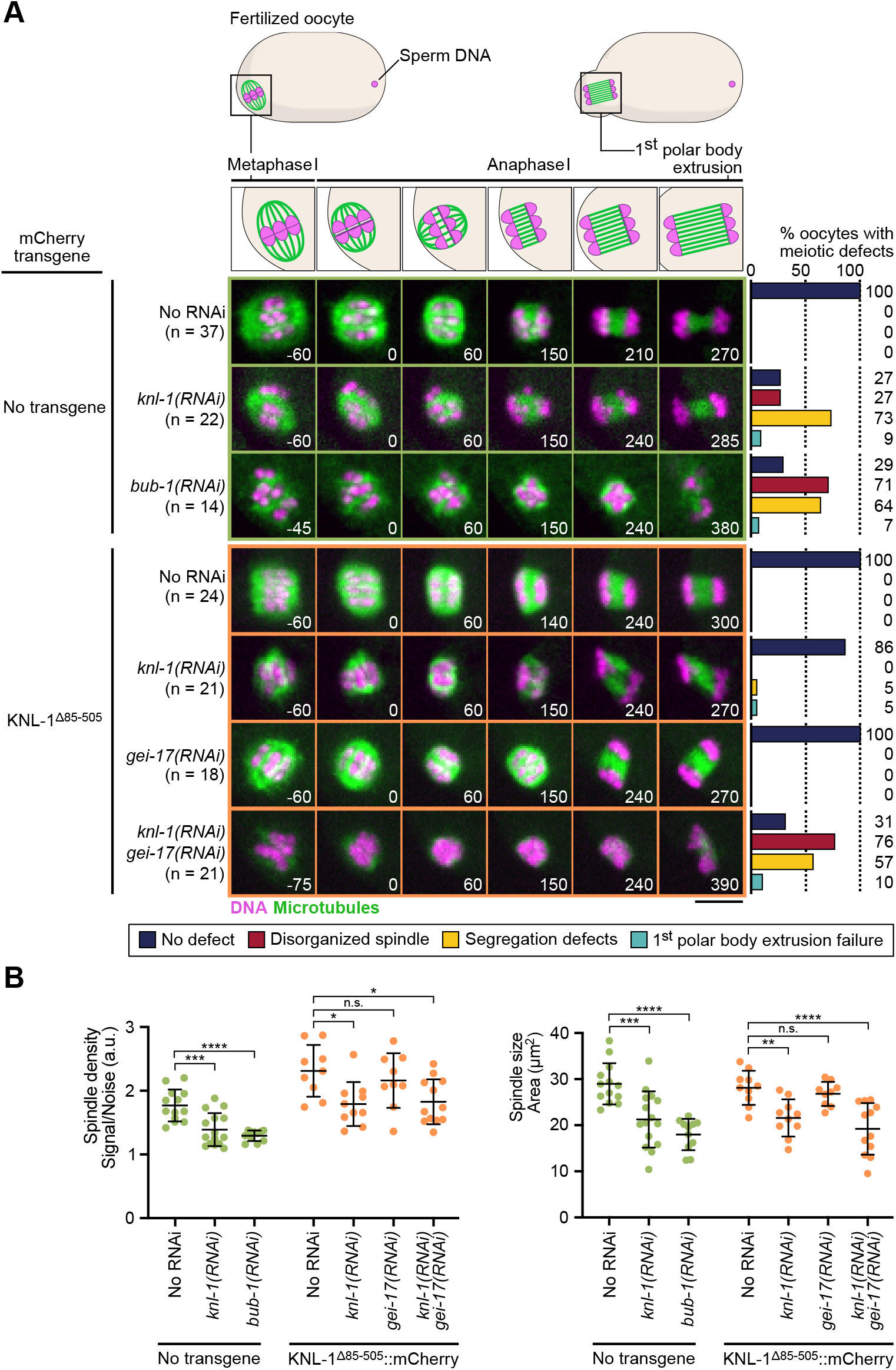
BHC module has both kinetochore-dependent and - independent functions in spindle assembly and chromosome segregation in oocytes. (A) Schematics of the meiotic spindle during meiosis I division (top) and stills from live imaging of meiosis I in indicated conditions (bottom). Microtubules (GFP::TBA-2^α-tubulin^) in green, chromosomes (mCherry::HIS-11^H2B^) in magenta. Time in seconds relative to anaphase I onset. Graphs on the right show quantifications of meiotic defects. Scale bar 5 µm. (B) Plots of spindle density (corrected GFP intensity, left) and spindle area (right) 45 seconds before anaphase I onset. One-way ANOVA multiple comparisons, alpha=0.05, *n.s.* not significant.

To determine if the mild spindle phenotype observed in KNL-1^Δ85-505^ mutant oocytes after KNL-1 depletion, is due to the persistence of the ring domain BHC pool, we depleted GEI-17 in these oocytes. Consistent with our hypothesis, the mild spindle phenotypes observed in the absence of BHC module localization at kinetochores was strongly aggravated upon simultaneous BHC delocalization at ring domains after simultaneous depletion of KNL-1 and GEI-17 in KNL-1^Δ85-505^ mutant oocytes (Figure 3A-B, Movie 2). Therefore, functions of kinetochore and ring domain pools of the BHC module are partially redundant in the control of spindle assembly and chromosome segregation. This finding also implies that CLS-2 is primarily acting from chromosomes and not as a diffuse pool of protein along the spindle.

### BHC module integrity is essential for spindle assembly and chromosome segregation (Figure 4)

During mitosis in *C. elegans*, HCP-1/2 have been proposed to primarily act as CLS-2 kinetochore-targeting proteins ^22^. We tested if this was also the case during meiosis by analyzing the effect of compromising BHC module integrity in oocytes. For this, we analyzed spindle assembly and chromosome segregation in oocytes expressing the BUB-1^ΔKD^ mutant in absence of endogenous BUB-1, and the HCP-1^ΔCTD^ or HCP-1^ΔNTD^ mutants in the *hcp-2Δ* strain following depletion of endogenous HCP-1. The three mutants should prevent BHC module assembly via different means. BUB-1^ΔKD^ blocks HCP-1/2 and CLS-2 recruitment to the kinetochores and ring domains, but does not disrupt HCP-1/2 and CLS-2 binding. HCP-1^ΔCTD^ binds CLS-2 and does not disrupt KNL-1 and BUB-1 binding, but does not localize to kinetochores. HCP-1^ΔNTD^ localizes to kinetochores, but blocks CLS-2 kinetochore recruitment (Figure 4A). Spindle assembly was severely disrupted in all mutants with a monopolar spindle phenotype and reduced microtubule density. Meiotic chromosome segregation occurred, although lagging chromosomes during anaphase and retracted polar bodies during telophase were evident (Figure 4B-C, Movie 3, Figure 4 – Figure supplement 1A). Mitotic chromosome segregation was also strongly perturbed in the HCP-1^ΔCTD^ or HCP-1^ΔNTD^ mutants in absence of endogenous HCP-1/2 proteins, with evident sister chromatid co-segregation (Figure 4 – Figure supplement 1B). Overall, expression of BUB-1^ΔKD^, HCP-1^ΔCTD^ or HCP-1^ΔNTD^ following depletion and/or deletion of BUB-1 and HCP-1/2 recapitulated the *bub-1(RNAi)* and *hcp-1/2(RNAi)* phenotypes, with disorganized spindles and chromosome segregation defects. These results suggest that the primary function of BUB-1 and HCP-1/2 occurs in the context of the BHC module.

**Figure 4.**
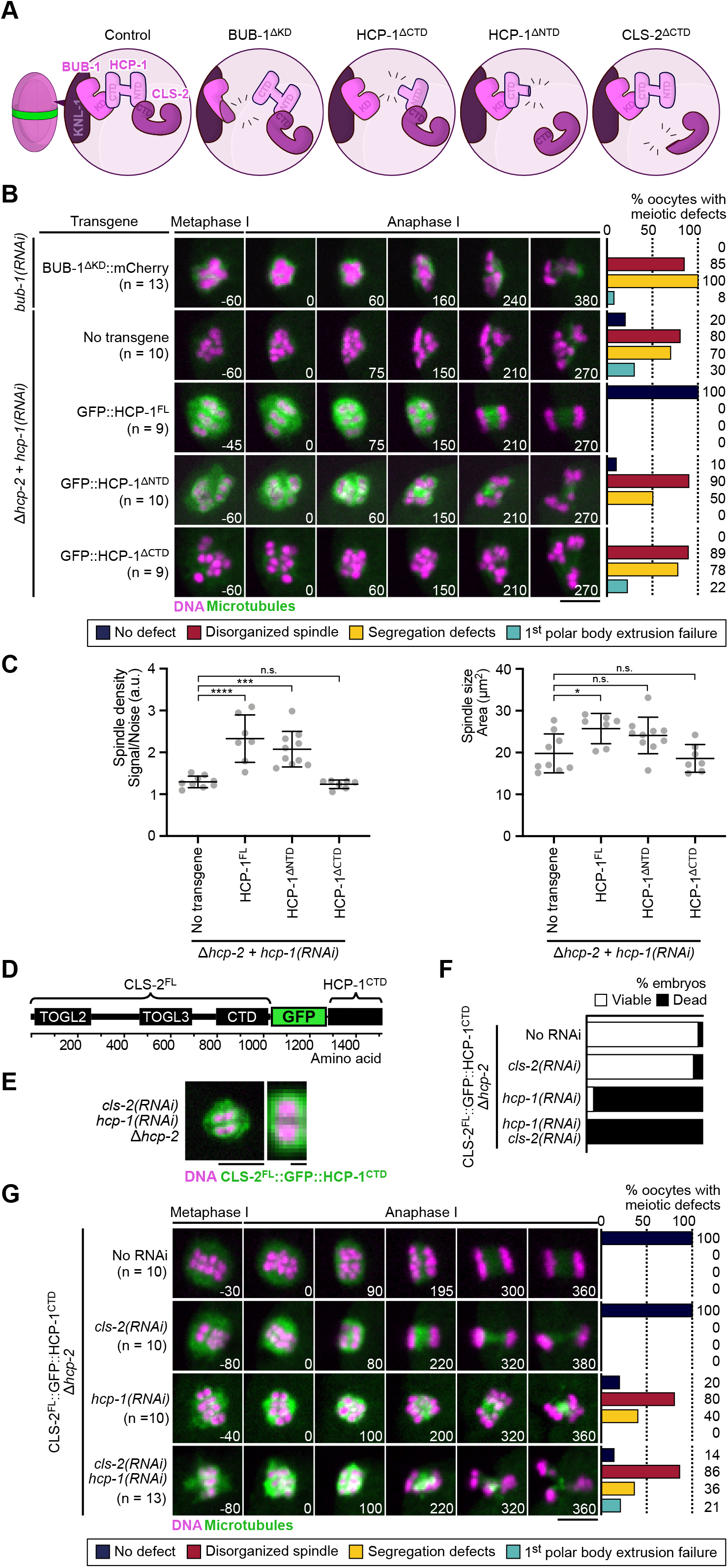
BHC module integrity is essential for spindle assembly and accurate chromosome segregation. (A) Schematics of BHC-module integrity mutants. (B) Stills from live imaging of meiosis I in indicated conditions. Microtubules (GFP::TBA-2^α-tubulin^) in green, chromosomes (mCherry::HIS-11^H2B^) in magenta. Time in seconds relative to anaphase I onset. Graphs indicate quantifications of meiotic defects. (C) Plots of spindle density (corrected GFP intensity, left) and spindle area (right) 45 seconds before anaphase I onset. Kruskal-Wallis multiple comparisons, alpha=0.05, *n.s.* not significant. (D) Schematics of the CLS-2^FL^::GFP::HCP-1^CTD^ protein fusion. (E) Localization of CLS-2^FL^::GFP::HCP-1^CTD^ (green) in metaphase I, and magnification of a single meiosis I chromosome. DNA (mCherry::HIS-11^H2B^) in magenta (n=13). (F) Embryonic lethality and (G) meiotic defects rescue assays by CLS-2^FL^::GFP::HCP-1^CTD^ in indicated conditions. Scale bars, full spindle 5 µm, single chromosome details 1 µm.

To further confirm that integrity of the BHC module is essential for its function, and that targeting CLS-2 to kinetochores is not sufficient for normal spindle assembly and chromosome segregation, we used protein engineering to allow CLS-2 kinetochore targeting in the absence of HCP-1/2. We engineered a fusion protein between GFP-tagged CLS-2 and the HCP-1 CTD (CLS-2::GFP::HCP-1^CTD^, Figure 4D), which we introduced in the *hcp-2Δ* strain. We then combined this engineered protein with *hcp-1(RNAi)* to artificially target CLS-2 to kinetochores independently of HCP-1/2. If BUB-1 and HCP-1/2 only act as CLS-2 adapter proteins, the CLS-2::GFP::HCP-1^CTD^ fusion should sustain embryonic development and viability in absence of HCP-1/2. We first verified that CLS-2 fused to GFP and to the HCP-1 CTD localized to kinetochores and to ring domains (Figure 4E), compared to GFP-tagged wild-type CLS2. Next to ensure that CLS-2::GFP::HCP-1^CTD^ was functional, we checked that it could sustain embryonic development and viability in absence of endogenous CLS-2 (Figure 4F). Accordingly, CLS-2::GFP::HCP-1^CTD^ also supported normal meiotic and mitotic spindle assembly and chromosome segregation upon *cls-2(RNAi)* in oocytes (Figure 4G, Movie 4). Therefore, fusing CLS-2 to GFP and to the HCP-1 CTD did not prevent its correct localization nor functionality. CLS-2::GFP::HCP-1^CTD^ also localized normally in absence of HCP-1 (Figure 4E), which shows that the HCP-1 CTD is both necessary and sufficient for localization. However, spindles and chromosome segregation in oocytes and zygotes were severely affected upon *hcp-1(RNAi)* (Figure 4G). Embryonic viability was also low in this condition (Figure 4F), suggesting that HCP-1/2 do not act only as targeting proteins for CLS-2 and likely have other roles in regulating microtubule dynamics in oocytes. We importantly obtained similar results when the GFP::HCP-1^CTD^ was fused to the CLS-2^ΔCTD^ transgene, which cannot interact with endogenous HCP-1/2 (Figure 4 – Figure supplement 1C-E, Movie 5). Overall, these results show that BHC module integrity and proper localization is essential for meiotic spindle assembly and chromosome segregation in oocytes.

### CLS-2 functions through a single TOGL domain (Figure 5)

We next focused on the functional domains of CLS-2. In most species, CLASP proteins are comprised of two to three ordered TOGL (Tumor Overexpressed Gene Like 1, 2 and 3) domains ^80^. In human CLASP2, these domains have specific functions. TOGL2 is essential for catastrophe suppression, whereas TOGL3 enhances rescues ^27, 28, 81^. The C-terminal domain (CTD) of CLASPs, responsible for kinetochore targeting, can inhibit these functions in human CLASP2, while the TOGL1 is required to release this auto-inhibition ^28^. Additionally, a conserved Serine/Arginine-rich (S/R-rich) region, important for microtubule lattice binding through electrostatic interactions, separates TOGL2 and 3 ^24, 82^. Sequence analysis showed that CLS-2 contains only two TOGL domains (TOGL2 and 3) separated by an S/R rich region, and a CTD separated from TOGL3 by a linker region predicted to be largely unfolded and a small also unfolded C-terminal tail (Figure 5 – Figure supplement 1A, B).

**Figure 5.**
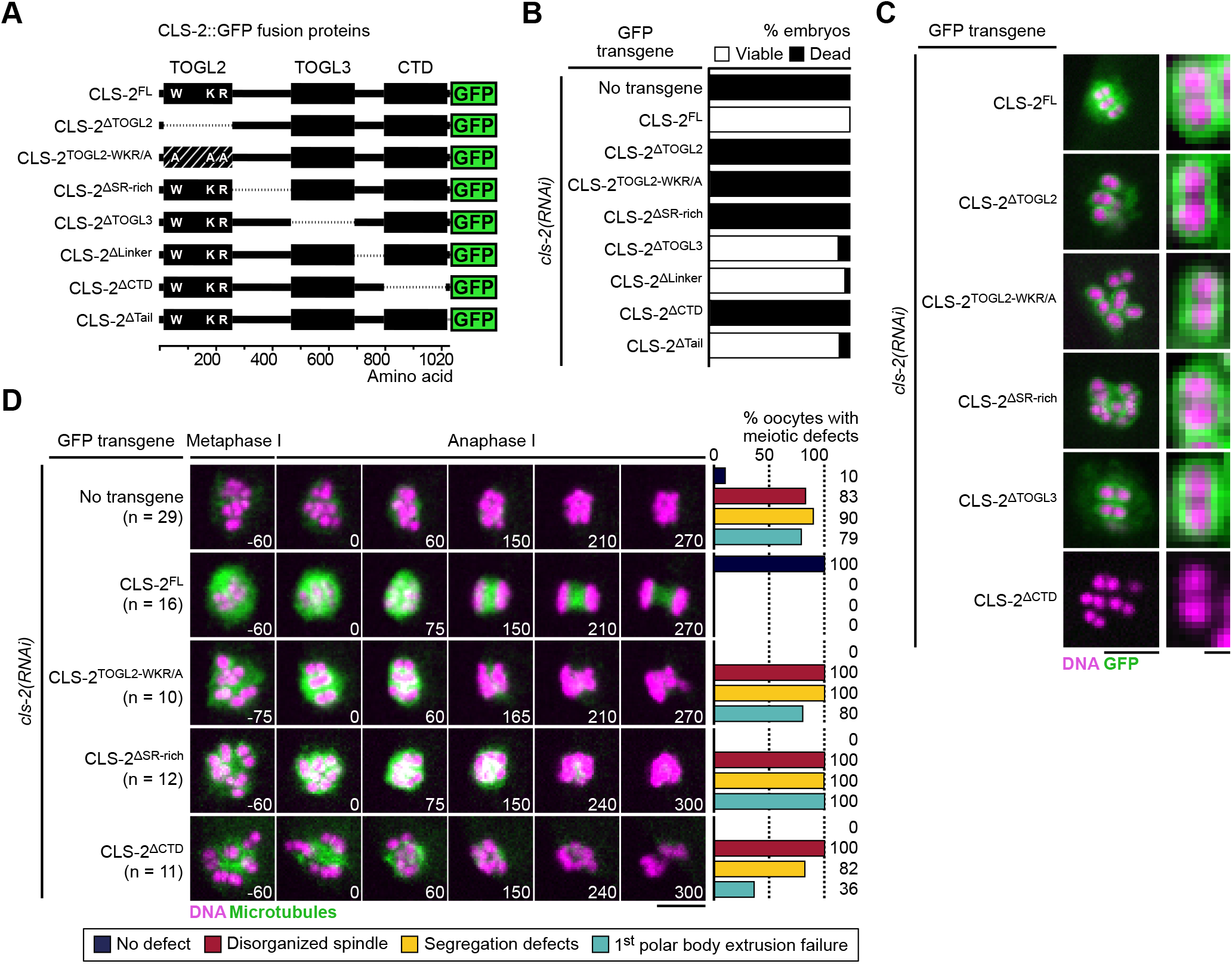
CLS-2 functions through a single TOGL domain. (A) Schematics of CLS-2::GFP truncated fusions. (B) Embryonic viability assay upon depletion of endogenous *cls-2* in the presence of indicated transgene. (C) Localization of CLS-2::GFP truncations (green) during metaphase I in indicated conditions. DNA (mCherry::HIS-11^H2B^) in magenta (n≥9). (D) Stills from live imaging of meiosis I. Microtubules (GFP::TBA-2^α-tubulin^) in green, chromosomes (mCherry::HIS-11^H2B^) in magenta. Time in seconds relative to anaphase I onset. Graphs indicate quantifications of meiotic defects. Scale bars, full spindles 5 µm, single chromosome details 1 µm.

We generated a series of transgenic strains expressing RNAi-resistant GFP-fused truncations in CLS-2 or point mutations in conserved and functionally important residues of TOGL2 (Figure 5A, Figure 5 – Figure supplement 1C), and we quantified embryonic viability upon depletion of endogenous CLS-2. All transgenes were expressed at comparable levels, but only four mutants were unable to sustain embryonic viability in absence of endogenous CLS-2. These corresponded to deletion or point mutations within TOGL2 (CLS-2^ΔTOGL2^, CLS-2^TOGL2-WKR/A^), the S/R-rich region (CLS-2^ΔSR-rich^), and the CTD (CLS-2^ΔCTD^) (Figure 5B). In contrast, deleting TOGL3, or the linker region between TOGL3 and the CTD, or the C-terminal tail had no significant effect on embryonic viability upon depletion of endogenous CLS-2 (Figure 5B). Sequence analysis revealed that TOGL3 lacks several conserved amino-acids (present in TOGL2 and mutated in the CLS-2^TOGL2-WKR/A^ mutant) corresponding to residues that normally contribute to microtubule binding in human CLASPs and to tubulin binding in XMAP215 family proteins ^83, 84^. This evolutionary sequence divergence could potentially account for the apparent lack of requirement for TOGL3, as mutating the corresponding residues in TOGL2 abrogated CLS-2 functionality (Figure 5B, D). As indicated previously, only deleting the CTD prevented chromosomal localization of the corresponding GFP-tagged transgene (Figure 5C), suggesting that the other three transgenes carried loss-of-function mutations.

We confirmed a loss-of-function phenotype by analyzing meiotic spindle assembly and chromosome segregation in oocytes lacking endogenous CLS-2. Mutating TOGL2, or deleting the SR-rich region or the CTD led to phenotypes comparable to the full loss of function of CLS-2 with severely disorganized spindles, inaccurate chromosome segregation and polar body extrusion failures (Figure 5D, Movie 6). While mutations in the TOGL2 or SR-rich region usually led to complete abrogation of chromosome segregation (70% and 92% oocytes did not attempt chromosome segregation, respectively), and to polar body extrusion failure (80% and 100% oocytes, respectively), oocytes lacking the CLS-2 CTD frequently displayed attempted chromosome segregation (only 18% oocytes did not attempt chromosome segregation) and often successfully extruded the first polar body (36% polar body extrusion failure). Chromosome segregation defects were also observed in mitosis in the presence of the corresponding transgenes upon depletion of endogenous CLS-2 (Figure 5 – Figure supplement 1D). Thus, as in human CLASP2, CLS-2 functions via a single TOGL domain (TOGL2), and also requires the CTD for HCP-1 binding and the S/R-rich region for potential microtubule lattice-interaction.

### BHC module components synergistically stabilize microtubules *in vitro* (Figure 6)

To investigate the effect of BHC module components on microtubule dynamics, we purified full-length BUB-1, HCP-1, and GFP-tagged CLS-2 from insect cells and analyzed their activity using an *in vitro* microtubule-based assay (Figure 6 – Figure supplement 1A). Microtubule growth from GMPCPP-stabilized seeds was monitored by TIRF (Total Internal Reflection Fluorescence) microscopy at 23°C ^85, 86^ (Figure 6A). Consistent with BUB-1 not being a microtubule-associated protein (MAP), analysis of microtubule dynamics showed that 100 nM BUB-1 did not have any effect on the microtubule growth rate, on catastrophe and rescue frequencies, nor on the time spent in pause by microtubules (Figure 6 B-F, Movie 7). BUB-1 on its own also did not interact with microtubules in a pelleting assay (Figure 6 – Figure supplement 1B), nor show any microtubule-bundling activity (Figure 6G). In contrast, both HCP-1 and GFP-CLS-2 independently displayed strong microtubule-bundling activity (Figure 6G). Consistent with a previous study showing that human CENP-F can stimulate microtubule polymerization *in vitro* ^62^, 100 nM HCP-1 had a mild but significant promoting effect on the microtubule growth rate (1.4-fold increase). This was interestingly abrogated by addition of 100 nM BUB-1, but not by a combination of BUB-1 and GFP-CLS-2 (100 nM each) (Figure 6 B-F, Movie 7). In contrast, 100 nM GFP-CLS-2 alone had the opposite effect and inhibited the microtubule growth rate (1.1-fold reduction), which is consistent with the effect of human or *Drosophila* CLASPs on microtubule dynamics ^25, 26, 28^. Also consistent with previous reports on CLASPs, we found that GFP-CLS-2 had significant catastrophe-suppressing, and rescue- and pause-promoting activities (Figure 6 B-F, Movie 7) ^24, 27, 28^. Furthermore, and like human CLASPs, CLS-2 is unlikely to function as a microtubule polymerase, as neither of the two TOGL domains interacted with free tubulin (Figure 6 – Figure supplement 1C) ^28^. Importantly, the effect of GFP-CLS-2 on catastrophe and rescue was independent of HCP-1 and/or BUB-1 (Figure 6C-F). In contrast, the most surprising effect of reconstituting the BHC module (BUB-1, HCP-1 and GFP-CLS-2 together) *in vitro* was the strong increase in the pause-promoting activity (31-fold increase in the time spent in pause) compared to all other conditions tested (Figure 6F, Movie 7). Thus, reconstituting the BHC module *in vitro* displays both additive (promotion of the growth rate and rescue frequency, and inhibition of catastrophe) and synergistic (promotion of pause) effects on microtubule dynamics compared to individual components, which leads to microtubule stabilization. This also indicates that BUB-1 and HCP-1 can modulate the effect of CLS-2 on microtubule dynamics. Overall, our *in vitro* results are consistent with our *in vivo* data, with BUB-1 and HCP-1/2 being important contributors to the regulation of microtubule dynamics, and not simply acting as CLS-2 targeting proteins, in the context of the BHC module.

**Figure 6.**
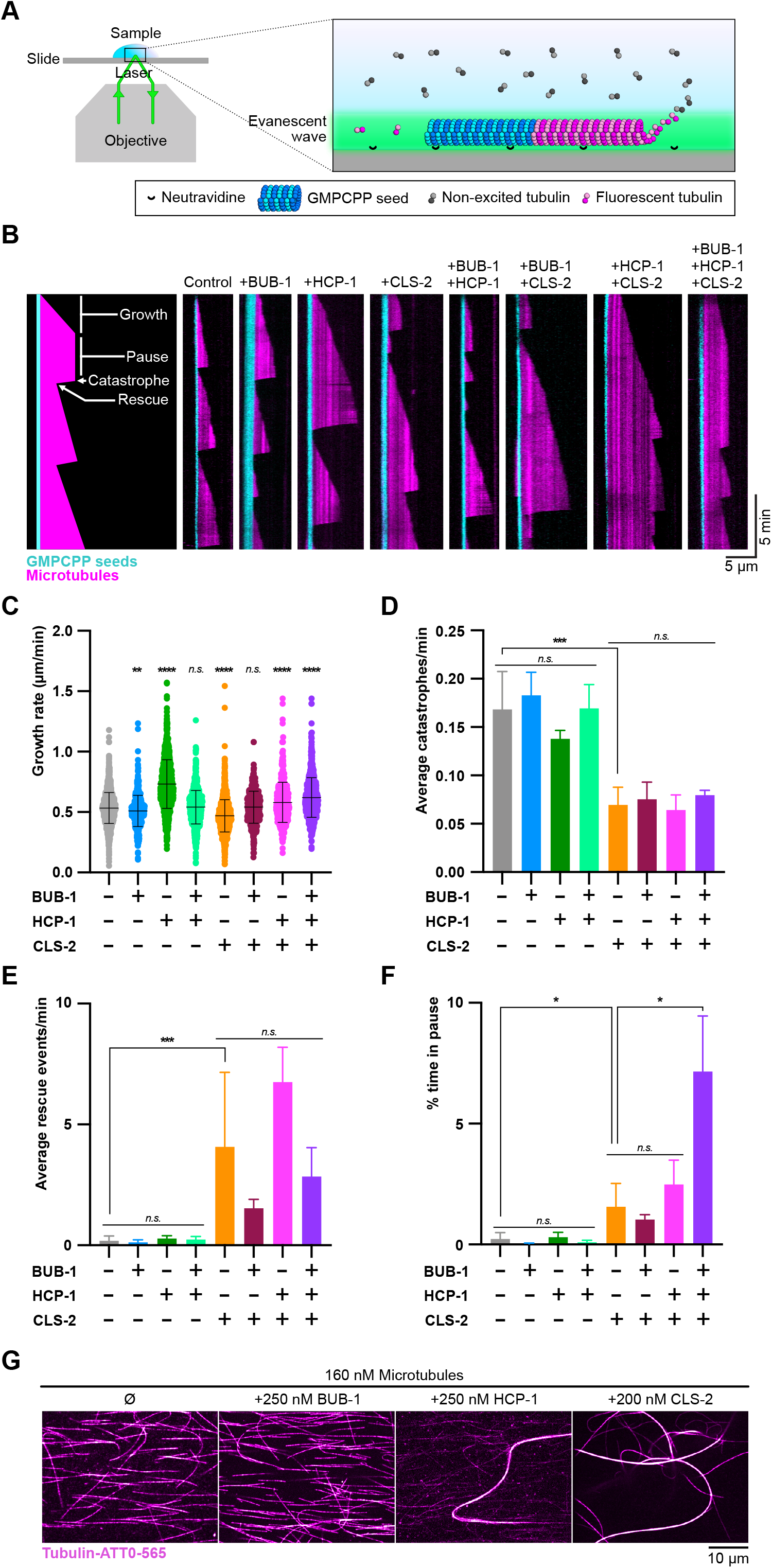
BHC module components synergistically stabilize microtubules *in vitro*. (A) Schematics of the TIRF microscopy-based microtubule assay. Labeled tubulin (ATTO-565, magenta) fluoresces only when at the surface of the coverslip. Microtubules polymerize from biotinylated GMPCPP seeds (tubulin-ATTO-488, cyan) bound to a Neutravidine-coated glass coverslip. (B) Representative kymographs of microtubules (magenta) growing from GMPCPP seeds (cyan) in the presence or absence of BUB-1, HCP-1 and/or CLS-2 (100 nM each). Schematics on the left highlights the different observed microtubule dynamics events. (C-F) Quantification of growth (C), catastrophe (D), rescue (F), and pause (D) events. One-way ANOVA, alpha=0.001 (C), and Kruskal-Wallis multiple comparison tests (D-F), alpha=0.05, *n.s.* not significant. (G) Microtubule bundling assay. Organization of microtubules (magenta) observed in indicated conditions.

## DISCUSSION

Our results highlight the importance and molecular mechanisms by which a kinetochore module, the BHC module, comprising the kinase BUB-1, the two CENP-F orthologs HCP-1/2 and the CLASP family member CLS-2, regulate microtubule dynamics in *C. elegans* oocytes and zygotes for efficient chromosome segregation.

### BHC module assembly and kinetochore targeting

We found that assembly of the BHC module requires the kinase domain of BUB-1 and the CTD of HCP-1, although we were unable to show a direct interaction between these domains using a yeast-two-hybrid assay. This may be a false negative due to proteins not being properly expressed/folded in yeast. Supporting this view, in humans, the kinase domain of Bub1 interacts directly with a C-terminal domain of CENP-F, and this interaction is necessary for CENP-F kinetochore targeting, which is reminiscent of our findings in *C. elegans* oocytes and zygotes ^46^. Also consistent with previous findings in human cells and with the fact that a kinase dead mutant of BUB-1 can sustain embryonic viability in *C. elegans*, we show that the kinase domain, but not kinase activity of BUB-1, is required for the localization of the CENP-F-like protein HCP-1 to kinetochores ^79^. Finally, we found that CLS-2 kinetochore recruitment involves a direct interaction between the HCP-1 NTD and CLS-2 CTD. In humans, CLASP kinetochore localization was also shown to depend on its CTD ^29^. However, in contrast to our present results, CENP-E, but not CENP-F, is responsible for kinetochore targeting of CLASP ^31^. Interestingly, CENP-E and CENP-F have been speculated to be distantly related paralogs ^46^. The fact that in *C. elegans*, which lack a CENP-E ortholog, the CENP-F-like protein HCP-1 is responsible for CLASP kinetochore localization supports this hypothesis.

A surprising finding was that kinetochore targeting of the BHC module could occur independently of BUB-3 in *C. elegans* oocytes, which is in contrast with previous findings in *S. cerevisiae* and human cells ^51, 87^. How then is BUB-1 targeted to kinetochores in *C. elegans*? We envision two plausible scenarios. First, although Bub3, in yeasts and mammals, is the primary determinant of Knl1 MELT binding, structural analysis demonstrated that direct contacts, which normally reinforce Bub3 binding, exist between a unique loop in Bub1 and the Knl1 MELT repeats ^88^. In *C. elegans*, these contacts could be sufficient for efficient BUB-1 recruitment to kinetochores in absence of BUB-3. Second, prior to the identification of the primary role of Knl1 MELT repeats for Bub1/Bub3 kinetochore targeting, a prevalent model implicated interaction between a KI (Lys-Ile) motif in Knl1, which lies in the MELT repeat region of the protein, and the conserved TPR domain of Bub1 ^89, 90^. Although, the KI motif was later shown to only act as a MELT-enhancing motif for Bub3-binding, and no clear equivalent motif could be identified in *C. elegans* KNL-1, it is possible that a sequence-divergent, but functionally equivalent, motif exists in KNL-1 that would be sufficient for recruiting BUB-1 to kinetochores ^91^. Furthermore, the fact that in *C. elegans* a *bub-3Δ* mutant is viable and fertile implies that it does not display any of the severe meiotic and mitotic phenotypes associated with loss-of-function of the BHC module ^78^. This in turn indicates that the reduced BHC pools, recruited at kinetochores and ring domains independently of BUB-3, are functional. Further functional analysis of BUB-1 kinetochore targeting in this system will be required to elucidate the BUB-3-independent mechanism of BHC module localization.

### BHC function in meiotic spindle assembly and chromosome segregation

Our findings demonstrate that integrity of the BHC module is essential *in vivo* in *C. elegans* oocytes for functional meiotic spindle assembly. Specifically, our quantitative analysis of meiotic spindle assembly and function, demonstrated that depletion of BHC components individually, or perturbation of BHC module assembly, gave rise to disorganized spindles with reduced microtubule densities. Although mammalian CLASPs have also been involved in proper spindle assembly and mitotic fidelity, this contrasts with previous work in mammals, which demonstrated that CENP-F is largely dispensable for spindle assembly and chromosome segregation ^72, 73, 92, 93^. While in mammals CENP-E, and not CENP-F, is responsible for localizing CLASPs at kinetochores, a critical function of *C. elegans* HCP-1/2 is to target the CLASP family member CLS-2 to kinetochores and ring domains ^31^. Yet, our *in vivo* results show that localizing CLS-2 to kinetochores and rings independently of HCP-1/2 is not sufficient to rescue the spindle phenotype of HCP-1/2 depleted oocytes. This argues for HCP-1/2 playing additional essential roles in regulating microtubule dynamics and spindle function, beside simply promoting CLS-2 localization.

In human cells, CENP-F binds to microtubules through two independent MTBDs located at either terminus of the protein, both of which are required for normal tension at centromeres in human cells ^62, 64, 65, 94^. Our *in vitro* experiments are consistent with HCP-1 also binding directly to microtubules, although the domain(s) responsible for this activity has not been identified. Human CENP-F was also shown to stimulate microtubule assembly in a bulk *in vitro* assay through an unknown mechanism ^62^. We interestingly show here that the *C. elegans* orthologous protein HCP-1 increases the microtubule growth rate *in vitro*. Whether microtubule binding and/or microtubule growth rate promotion, independent of CLS-2 targeting, are the important functions of HCP-1 explaining its direct role in meiotic spindle assembly is unclear. Further work, including identification of separation-of-function mutations of HCP-1, is required to establish the CLS-2-independent role of HCP-1 *in vivo*.

Our data also show that the CTD deletion mutant of CLS-2 (CLS-2^ΔCTD^), which was unable to interact with HCP-1 or support normal meiotic spindle assembly, could largely support chromosome segregation and polar body extrusion, unlike other CLS-2 loss-of-function mutants. This suggests that CLS-2^ΔCTD^ has retained some functions of CLS-2, and extends previous findings showing that CLS-2 can localize to, and promote assembly of, central spindle microtubules in absence of its upstream partner protein BUB-1, although less efficiently ^35^. Our *in vitro* microtubule-based assays demonstrated that as a complex the BHC module promotes microtubule rescue, reduces catastrophe, and strongly enhances microtubule pausing relative to individual BHC module components individually or in pairs. Altogether, our data suggest that meiotic spindle assembly, chromosome segregation, and polar body extrusion are distinct functions of CLS-2 requiring different levels of microtubule dynamics regulation. Meiotic spindle assembly requires an intact BHC module and the strong microtubule pause promoting effect of the module, while chromosome segregation and polar body extrusion only depend on the rescue promoting and catastrophe reducing effects of CLS-2 alone.

In mammals, CLASPs contain two tandemly-arranged SxIP motifs that mediate End-Binding (EB) protein interaction important for microtubule plus-end accumulation and microtubule catastrophe suppression ^28, 81, 82, 95^. In *C. elegans* oocytes and embryos, CLS-2 did not substantially accumulate at microtubule plus-ends, which was consistent with the apparent lack of SxIP motif in the CLS-2 sequence. We noted that CLS-2 contained a divergent LxxPTPh motif (LPKRPTPQ) at the C-terminal end of the SR-rich region, which could potentially mediate EB interaction ^96^. However, a GFP-fused deletion of the corresponding region could support embryonic viability in absence of endogenous CLS-2, which suggests that it is not essential (Figure 6 – Figure supplement 1D). Also, in contrast to human CLASP2, which weakly but visibly accumulates at microtubule plus-ends *in vitro* in the absence of EB protein, GFP-tagged CLS-2 only faintly decorated the microtubule lattice, with a speckled pattern (Figure 6 – Figure supplement 1E). This weak microtubule binding was not increased in the presence of HCP-1 and BUB-1 (Figure 6 – Figure supplement 1E). Finally, in contrast to work on yeasts and *Drosophila* CLASPs, but in line with previous *in vitro* studies on human CLASPs, we did not detect any significant correlation between the CLS-2 speckles along the lattice and the sites of microtubule rescue or pause events (Figure 6 – Figure supplement 1F) ^24, 26, 27^. Therefore, unlike mammalian, or yeasts and *Drosophila* CLASPs, which respectively prevent catastrophe and promote rescue by concentrating at a region located behind the outmost microtubule end, or by accumulating at speckles along the lattice, the stabilizing effect of *C. elegans* CLS-2 does not require its plus-end nor lattice speckled accumulation. Instead binding to HCP-1 and BUB-1 could modulate CLS-2 effect on microtubules. It would thus be interesting to determine if similar synergistic regulation of microtubule dynamics exists between other CLASP proteins and their multiple targeting proteins. In particular, determining if CLASPs kinetochore-targeting by CENP-E and Astrin can also modulate their microtubule dynamics-regulating activity will be essential to fully understand CLASPs function in the control of chromosome segregation in mammals.

## METHODS

### *C. elegans* strain maintenance

*C. elegans* strains were maintained at 20° or 23°C under standard growth conditions on NGM plates and fed with OP50 *E. coli* ^97^. The N2 Bristol strain was used as the wild-type control background ^98^, unless specified otherwise in the text. Transgenic lines were obtained either by Mos1-mediated Single Copy Insertion (MoSCI) ^99^, or by crossing pre-existing strains. The list of strains used in this study is provided in Supplementary Material, Table S1. All strains produced will be provided upon request and/or made available through the Caenorhabditis Genetics Center (CGC, https://cgc.umn.edu/).

### RNA-mediated interference

Double-stranded RNAs were produced as described previously in ^39^. After DNA amplification from total N2 cDNA by PCR using the primers listed below, reactions were cleaned (PCR purification kit, Qiagen), and used as templates for T3 and T7 transcription reactions (MEGAscript,Invitrogen) for 5 hr at 37 °C. These reactions were purified (MEGAclear kit, Invitrogen), then annealed at 68 °C for 10 minutes, 37 °C for 30 minutes. L4-stage hermaphrodite larvae were micro-injected with 1500-2000 µg/µL of each dsRNA, and recovered at 20°C for 36 hours before being further processed.

Primers used to produce double-stranded RNAs targeting the indicated genes:

*bub-1* (R06C7.8):

5‘-AATTAACCCTCACTAAAGGggataattttatgatcaccag-3’

5’-TAATACGACTCACTATAGGctacttttggttggcggcaag-3’

*cls-2* (R107.6):

5’-TAATACGACTCACTATAGGttcaaggaaaagttggacc-3’

5’-AATTAACCCTCACTAAAGGggtgcatttctgattccacc-3’

*gei-17* (W10D5.3):

5’-AATTAACCCTCACTAAAGGTATGCTGATAATTTTGAACCGCT-3’

5‘-TAATACGACTCACTATAGGTCATCAACAATAAGTCTATCATATGG-3’

*hcp-1* (ZK1055.1):

5‘-AATTAACCCTCACTAAAGGaagcgccagcaaaccgagtcgcc-3’

5’-TAATACGACTCACTATAGGgtcaatgtgacctttgacaggaagc-3’

*knl-1* (C02F5.1):

5’-TAATACGACTCACTATAGGttcacaaacttggaagccgctg-3’

5’-TAATACGACTCACTATAGGttcacaaacttggaagccgctg-3’

### Embryonic viability assays

Embryonic viability assays were performed at 20°C. Worms were singled onto plates 36h post-L4, upon recovery from dsRNA micro-injection. Worms were allowed to lay eggs for 12h before being removed from the laying plates. For each laying plate, the number of unhatched eggs, and of L1 larvae was counted. Plates were then left at 20°C for another 36h before the total number of worms reaching L4/adulthood was counted. Control worms were analyzed with the same protocol with no preceding micro-injection. The proportion of viable progenies was calculated as the number of L4/adults divided by the number of eggs/L1.

### Live Imaging and image analysis

All live and fixed acquisitions were performed on a Nikon Ti-E inverted microscope, equipped with a Yokogawa CSU-X1 (Yokogawa) spinning-disk confocal head with an emission filter wheel, using a Photometrics Scientific CoolSNAP HQ2 CCD camera. The power of 100 or 150 mW lasers was measured before each experiment with an Ophir VEGA Laser and energy meter. Fine stage control was ensured by a PZ-2000 XYZ Piezo-driven motor from Applied Scientific Instrumentation (ASI). The microscope was controlled with Metamorph 7 software (Molecular Devices).

For *ex utero* live imaging of embryos and one-cell zygotes, embryos were freed by dissecting worms on a cover slip in 6-8 µL meiosis medium ^100^. Movies were acquired using a Nikon APO λS x60/1.40 oil objective and 2×2 binning. Images were acquired at 10, 15 or 20 second interval, over 2 to 4 Z-planes with a step size of 2 µm. Temperature was maintained at 23°C using the CherryTemp controller system (Cherry Biotech).

Immunofluorescence was performed as described in ^101^ using a 20 minutes cold (−20°C) methanol fixation. Dylight 550-labeled rabbit anti-BUB-1 (custom made, ^74^) and FITC-labeled mouse anti-α-tubulin (SIGMA F2168, DM1α) were used at a concentration of 1 μg/mL. DNA was stained with Hoechst at 2 μg/mL. Z-sections were acquired every 0.2 μm using a Nikon APO λ x100/1.45 oil objective. Maximum projections of relevant sections are presented.

Image and movie treatment, scaling and analysis were performed in FIJI ^102^. Spindle area and fluorescence intensity were measured on sum projections of 4 Z-planes. The surface and microtubule density of the spindle were measured 45 seconds before anaphase I onset. For each sample, the background signal was measured using the same ROI placed at the center of the oocyte. Graphs represent ratios of the Integrated fluorescence/Background noise.

### Western blotting

For each sample, 50 gravid adult worms (24h post-L4) were washed in M9 buffer (22 mM KH_2_PO_4_, 42 mM Na_2_HPO_4_, 86 mM NaCl, and 1 mM MgSO_4_•7H_2_O) supplemented with 0,1% Triton X100. Samples were then resuspended in 30µL Laemmli buffer (1X final) and incubated at 97°C for 15 minutes, vortexed at 4°C for 15 minutes, and boiled at 97°C for 5 minutes. Worm extracts were loaded on NuPAGE 3-8% Tris Acetate gels (Invitrogen). Proteins were then transferred onto nitro-cellulose membranes which were incubated with 1 µg/µL primary antibodies in 1X TBS-Tween (TBS-T) supplemented with 5% skim milk. Anti-HCP-1 and anti-CLS-2 were custom-produced in rabbit ^74^. Mouse Anti-GFP (Roche 11814460001) was used to specifically detect the transgenes. Mouse anti-α-tubulin (Abcam Ab7291, DM1α) or rabbit custom-made anti-KLP-7 antibodies ^101^, were used as loading controls. Blockings were performed in 1X TBS-Tween+5% skim milk, and washes in 1X TBS-T. Signals were revealed with HRP-coupled goat anti-mouse or anti-rabbit secondary antibodies (Jackson ImmunoResearch 115-035-003 or 11-035-003, respectively, 1:10000 in 5% skim milk TBS-T). Revelation was performed on a Biorad ChemiDoc Imaging System.

### Yeast two hybrid assay

Full CDS or gene domains were cloned from wild type *C. elegans* (N2) cDNA. The yeast two hybrid assay was performed using the LexA-based system. In short, the HCP-1 NTD and CTD were fused to LexA binding domain in the bait pB27 plasmid, and other CDS sequences fused to Gal4 activating domain in the prey pP6 vector. Bait-encoding plasmids were transformed into L40Δga14 *S. cerevisiae* strain (mat α), prey plasmids into CG1945 strain (Mat a). Upon mating, diploid *S. cerevisae* were spotted on −Leu −Trp double-selection medium, and interactions were tested on −Leu −Trp −His medium. 5 mM of 3-amino-1,2,4-triazole (3AT) was added to abolish autoactivation by LexA::HCP-1(NTD). Image acquisitions were done on a Biorad ChemiDoc Imaging System.

### *In silico* protein sequence analyses

For comparison of TOGL domains among CLASPs (Ce: *Caenorhabditis elegans*, Dm: *Drosophila melanogaster*, Hs: *Homo sapiens*, Mm: *Mus musculus*, Xl: *Xenopus laevis*), protein sequence alignment and phylogenetic tree were generated using Clustal Omega (Version 1.2.4, https://www.ebi.ac.uk/Tools/msa/). The tree was rooted using the sequence of the TOG3 of human chTOG. The phylogenetic tree figure was generated using the FigTree software (v1.4.4, http://tree.bio.ed.ac.uk/software/figtree/).

Sequence alignment of the TOGL2 domains (At: *Arabidopsis thaliana*, Ce: *Caenorhabditis elegans*, Dm: *Drosophila melanogaster*, Hs: *Homo sapiens*, Mm: *Mus musculus*, Xl: *Xenopus laevis*) and the TOGL3 domain of *C. elegans* was done with MAFFT using the Snapgene software (https://www.snapgene.com/).

### Protein production and purification

Full length BUB-1 and HCP-1 sequences were amplified from *C. elegans* cDNA and cloned in pFastBac dual expression vector in frame with a 6xHis tag. CLS-2 was cloned in pFastBac dual in frame with a C-terminal GFP tag, aTEV protease cleavage site, and a 6xHis tag. BUB-1 and CLS-2::GFP production was performed in 2 L SF9 cells in Insect-XPRESS (Lonza) medium (1×10^6 cells/mL), infected with amplified baculovirus for 48h at 27°C. HCP-1 was produced similarly but in Hi-5 cells maintained in EX-Cell 405 medium (Sigma-Aldrich) and collected after 66h infection. Cells were harvested by centrifugation at 700xg and resuspended in 10-15mL lysis buffer (150 mM KCl, 1mM MgCl_2_, 0.1% Tween-20, protease inhibitor (Complete EDTA-free tablets, Roche)) per liter of culture, supplemented with 5% glycerol for CLS-2::GFP. BUB-1 was purified in 25 mM HEPES pH 7.2, HCP-1 in 50 mM MES pH6.5, and CLS-2::GFP in 50 mM PIPES pH 6.8 supplemented with 50 mM glutamate and 50 mM arginine to increase protein solubility and purification yield ^27, 103^. Cells were homogenized in a Dounce homogenizer and lysed by sonication for 30 seconds at 50% amplitude using a 6 mm diameter probe. Lysates were clarified by ultracentrifugation at 100,000xg for 1h at 4°C. Sample loading, column washes and elutions were performed using an ÄKTA™ Pure chromatography system (Cytiva). Supernatants were loaded on 1 mL HiTrap™ TALON™ crude (Cytiva) columns equilibrated with Buffer A (150 mM KCl, 1 mM MgCl_2_). The columns were washed with 10 column volumes (CVs) of buffer A followed by 30 CVs of buffer A’ (150 mM KCl, 1 M MgCl_2_). Columns were then equilibrated with 20 CVs of buffer A. Elution was performed with 30 CVs of a gradient from 0 to 100% buffer B (150 mM KCl, 1 mM MgCl_2_, 300 mM Imidazole). A, A’ and B buffer pHs were adjusted to protein-specific lysis buffer pH. Absorbance was measured at 280 nm. Fractions were analyzed by SDS-PAGE and Coomassie staining. Fractions containing pure proteins were pooled, concentrated using Amicon Ultra-15 centrifugation units and desalted against buffer A using Econo-Pac® 10DG Desalting columns (Bio-Rad). Proteins were aliquoted, snap frozen in liquid nitrogen and stored at −80°C. CLS-2::GFP protein was further purified using a gel filtration column Superdex® 200 Increase 10/300 GL (Cytiva). Equilibration, loading and isocratic elution were made at a flow rate of 0.4 mL/min with 25 mM HEPES pH 7.2, 150 mM KCl, 1 mM EGTA, 1 mM MgCl_2_.

CLS-2 TOGL2 (aa1-276) and TOGL3 (aa441-699) domains were expressed in *E. coli* Rosetta2 cells using pProEx-HTb 6xHis expression vectors. Transformed bacteria were allowed to reach 0.4 OD600nm at 37°C. The cultures were then transferred to 20°C and expression was induced overnight (~18 hours) with 0.1 mM IPTG. Bacteria were harvested by centrifugation at 4000xg for 20 minutes resuspended and washed with 1xPBS by centrifugation. Cells were resuspended in lysis buffer (25 mM MOPS pH 7.2, 300 mM NaCl, 10 mM ß-mercaptoethanol) supplemented with protease inhibitors (Complete EDTA-free tablets, Roche). Cells were lysed by sonication using a 6 mm diameter probe at 50% amplitude for 2 minutes. The lysates were clarified by ultracentrifugation at 90,000xg for 1 hour at 4°C and loaded on 1 mL HisTrap Excel™ columns (Cytiva). Purification of CLS-2 domains was performed using the same procedure as BUB-1, HCP-1 and CLS-2::GFP (see previous paragraph) but using different composition for buffer A (25 mM MOPS pH 7.2, 300 mM NaCl, 10 mM ß-mercaptoethanol), A’ (25 mM MOPS pH 7.2, 1 M NaCl, 10 mM ß-mercaptoethanol) and B (25 mM MOPS pH7.2, 300 mM NaCl, 10 mM ß-mercaptoethanol, 300 mM imidazole). CLS-2 domains were dialyzed against 25 mM MOPS pH 7.4, 1 mM EGTA, 300 mM KCl, 10 mM ß-mercaptoethanol, frozen in liquid nitrogen and stored at −80°C.

Tubulin was purified from pig brains following high salt protocol and cycles of polymerization and depolymerization as in ^104^. Tubulin was then labeled with either NHS-ester-ATTO 565 (ATTO-TEC), NHS-ester-ATTO 488 (ATTO-TEC) or EZ-Link Sulfo NHS-LC-LC-Biotin (ThermoFisher #21338). Labeling dyes or linkers were removed by two cycles of polymerization/depolymerization ^105^. In brief, unlabeled polymerized tubulin was incubated 40 minutes at 37°C in the presence of 5 mM of succinimidyl ester-coupled reagent in labeling buffer (0.1 M HEPES pH 8.6, 1 mM MgCl_2_, 1 mM EGTA, 40% glycerol (volume/volume)). Microtubules were then spun down through a low pH cushion (60% glycerol, 1xBRB (80 mM K-PIPES pH 6.8, 1 mM MgCl_2_, 1 mM EGTA)), resuspended in 50 mM K-glutamate pH 7.0, 0.5 mM MgCl_2_, and left to depolymerized on ice for 30 minutes in a small glass Dounce homogenizer. Depolymerized labeled tubulin was recovered from a 120,000xg centrifugation at 2°C and resuspended in 4 mM MgCl_2_, 1 mM GTP, 1xBRB. An additional cycle of polymerization was performed at 37°C for 40 minutes. Microtubules were sedimented at 120,000xg for 20 minutes at 37°C and depolymerized in ice-cold 1x BRB buffer. Soluble tubulin was recovered from a 10 minutes 150,000xg centrifugation at 2°C, diluted to 15-20 mg/ml in 1xBRB, aliquoted, frozen in liquid nitrogen and stored at −80°C.

### Protein-protein interactions by size exclusion chromatography

To assess the interaction of the CLS-2 domains with soluble tubulin, an equimolar mix was made at a concentration of 25 µM of each protein in equilibration buffer (25 mM HEPES pH 7.0, 80 mM KCl, 1 mM EGTA, 1 mM MgCl_2_, 5% glycerol). Samples (50 µL) were loaded on a Superdex® 200 Increase 10/300 GL (Cytiva) at a flow rate of 0.4 mL/min. Proteins were followed by measuring the absorbance at 280 nm and peaking fractions were loaded and analyzed by SDS-PAGE and Coomassie staining.

### Microtubules bundling assays

A 100 µL mixture of 80 µM unlabeled and ATTO-565-labelled tubulin (12:1 ratio) was incubated 5 minutes at 4°C and centrifuged at 100,000xg for 10 minutes to remove aggregated labeled tubulin. The supernatant was left to polymerize at 35°C for 30 minutes in 1 mM GTP, 1xBRB buffer for 30 minutes. An equal volume of 1xBRB buffer with 20 µM docetaxel (Sigma-Aldrich) was added, and the reaction was further incubated for 15 minutes at room temperature before being centrifuged at 50,000xg for 10 minutes at 25°C. The supernatant was discarded, the pellet was gently washed with a volume of warm (35°C) 10 µM docetaxel, 1xBRB. The pellet was rehydrated by incubating for 10 minutes at room temperature in a volume of 10 µM docetaxel, and then resuspended by pipetting up and down. The stabilized microtubule solution was diluted in 10 µM docetaxel,1xBRB to a final tubulin concentration of ~160 nM to facilitate visualization under the fluorescent microscope. The diluted microtubule suspension was incubated 5 minutes at room temperature with the protein of interest at a concentration of 200 nM to 250 nM in a microtube. The mixture was transferred in a ~10 µL microchamber between a microscope slide and a coverslip assembled with thin strips of double-sided tape. The chamber was immediately imaged with a spinning-disk confocal. An image of a single focal plane was captured at 60x magnification with a 1.4 N.A. oil immersion objective, using 561 nm excitation laser.

### Microtubules pelleting assays

Stabilized microtubules were prepared as stated above except that no labeled tubulin was added. Final reaction volumes of 50 to 100 µL of 1 µM purified BUB-1 and 0 to 1.7 µM stabilized microtubules were prepared in 10 µM docetaxel, 80 mM KCl, 1x BRB buffer, into ultracentrifugation microtubes. Mixtures were incubated 15 minutes at room temperature and centrifuged at 50,000xg for 10 minutes. The supernatants were kept in 1x Laemli sample buffer (LSB) from a 5x solution (400 mM TRIS-HCl pH6.8, 450 mM DTT, 10% SDS, 50% glycerol, 0.006% w/vol bromophenol blue). Pellets were washed with 10 µM docetaxel,1xBRB and directly resuspended in 1x LSB. Supernatants and pellets were analyzed by SDS-PAGE (10% acrylamide) and Coomassie staining.

### *In vitro* microtubule dynamics

Biotinylated GMPCPP-stabilized microtubule seeds were obtained by mixing biotinylated-tubulin and fluorescent tubulin (ATTO488-tubulin) at a 4:1 ration to a 10 µM final concentration of tubulin. This mix was incubated at 37°C in 1x BRB supplemented with 0.5 µM GMPCPP (Jenabioscience). After an hour of incubation, docetaxel (Sigma-Aldrich) was added to a final concentration of 1 µM and the reaction was incubated for 30 minutes at 30°C. Microtubule seeds were pelleted at 100,000g for 10 minutes at 25°C. The pellet was resuspended in 1x BRB, 0.5 mM GMPCPP, 1 µM docetaxel. Seeds were aliquoted, frozen in liquid nitrogen and stored at −80°C in cryotubes for up to 3 weeks.

Single microtubule dynamics assays were performed in a ~20 µL flow chamber between a glass slide and a coverslip assembled with double-sided sticky tape. Glass slides were cleaned and passivated using a protocol adapted from ^85^. In brief, slides were washed successively in water, acetone, ethanol, 2% Hellmanex detergent for at least 30 minutes in each reagent in a glass beaker immersed into an ultrasonic bath. Glass slides were then treated with a 1 mg/mL PEG-silane solution (MW 30K; Creative PEG works) in 96% ethanol, 0.1% HCl. In addition, coverslips were plasma-cleaned for 2 minutes before being treated with PEG-silane-biotin (MW 10K Creative PEG works) overnight at room temperature under gentle agitation. Slides and coverslips were abundantly washed with large volumes of MilliQ-water and dried using pressurized air blowing. Dried slides and coverslips were stored at 4°C for a maximum of 3 weeks in clean plastic boxes. Prior to use, a microchamber of around 10 µL in volume was fabricated using a slide, a coverslip and double-sided tape (~70 µm thickness). 100 µL of 50 µg/mL neutravidin (Invitrogen A2666), BRB-BSA (1xBRB, 0.2% BSA) solution were flowed into the chamber. The chamber was then kept at room temperature (22-23°C) and never allowed to dry. The neutravidin solution was incubated for 5 minutes in the chamber. The chamber was washed twice for 1 minute with 100 µL of 0.1 mg/mL PLL-PEG (PLL20k-G35-PEG2k, Jenkem), and a third time with 300-400 µL of BRB-BSA. A solution of GMPCPP-stabilized microtubule seeds diluted in BRB-BSA was flowed into the chamber and incubated for 3 minutes. Excess of seeds was then removed by several washes of >300 µL BRB-BSA before introduction of the elongation mix containing 12 µM total tubulin (94% unlabeled pig brain tubulin, 6% labeled pig brain tubulin), in 40 mM PIPES pH6.8, 10 mM HEPES pH 7.5, 44 mM KCl, 5 mM MgCl_2_, 1.5 mM EGTA, 0.2% methylcellulose (1500 cP), 4 mM DTT, 1 mM GTP, 0.5 mM ATP buffer supplemented with 128 nM catalase, 500 nM glucose oxidase and 40 mM glucose antifading agents, and in the presence of 100 nM freshly thawed BUB-1, HCP-1 and/or CLS-2::GFP. Microtubule dynamics was monitored between 22.5°C and 23°C on an azimuthal TIRF microscope (Nikon Eclipse Ti2 equipped with the Ilas2 module, Gataca systems) using a 60x, 1.47NA oil immersion TIRF objective and a Photometrics Prime BSI sCMOS camera. Two-channel acquisitions (488 nm and 561 nm) were performed every 3 seconds for 20 minutes. Imaged were acquired at 50 ms exposure.

### Figure preparation, graphs and statistical analyses

Figures and illustrations were done in the Affinity Designer software (ver. 1.10.3). Graphical representation of data and statistical analyses were performed using the GraphPad Prism software (ver. 8.4.3 (471)). Statistical tests used are specified in the corresponding figure legends.

## ACKNOWLEDGEMENTS

We thank all members of the Pintard and Dumont labs for support and advice. We are grateful to Patricia Moussounda, Téo Bitaille and Vincent Maupu-Massamba for providing technical support. We thank Dhanya Cheerambathur for worm strains and Benoit Palancade for yeast two hybrid strains. We thank Jérémie Gaillard and members of the CytoMorphoLab for technical advices and assistance with TIRF microscopy and microtubule-based assays. Some strains were provided by the CGC, which is funded by NIH Office of Research Infrastructure Programs (P40 OD010440). N. Macaisne was supported by an ANR contract (grant ANR-19-CE13-0015), L. Bellutti was supported by a post-doctoral fellowship from the Fondation pour la Recherche Médicale (FRM). This work was supported by CNRS and University Paris Cité, by NIH R01GM117407 and R01GM130764 (JC. Canman), and by grants from the European Research Council consolidator grant (ERC-CoG) ChromoSOMe grant 819179 and from the Agence Nationale de la Recherche ANR-19-CE13-0015 (J. Dumont).

## AUTHOR CONTRIBUTIONS

N. Macaisne, L. Bellutti, K. Laband, F. Edwards, B. Lacroix and J. Dumont conceived of the project. L. Pitayu conducted pilot experiments and analysis on BUB-1; B. Lacroix and A. Gervais performed all *in vitro* experiments. N. Macaisne, L. Bellutti, K. Laband, F. Edwards conducted all other experiments and data analysis. T. Ganeswaran and H. Geoffroy provided technical support. J. Dumont and B. Lacroix designed all experiments. N. Macaisne, L. Bellutti, K. Laband, F. Edwards, G. Maton, J.C. Canman, B. Lacroix and J. Dumont made intellectual contributions and wrote the manuscript. N. Macaisne and J. Dumont made the figures.

## SUPPLEMENTAL MOVIE LEGENDS

**Movie 1**

Live imaging of meiosis I in indicated conditions. Microtubules (GFP::TBA-2^α-tubulin^) in green, DNA (mCherry::HIS-11^H2B^) in magenta. Time in seconds relative to anaphase I onset. Scale bar 5µm.

**Movie 2**

Live imaging of meiosis I in *knl-1Δ85-505::mCherry* transgenic worms in indicated conditions. Microtubules (GFP::TBA-2^α-tubulin^) in green, DNA (mCherry::HIS-11^H2B^) in magenta. Time in seconds relative to anaphase I onset. Scale bar 5µm.

**Movie 3**

Live imaging of meiosis I in indicated transgenic *hcp-2*-mutant worms upon depletion of endogenous *hcp-1*. Microtubules (GFP::TBA-2^α-tubulin^) in green, DNA (mCherry::HIS-11^H2B^) in magenta. Time in seconds relative to anaphase I onset. Scale bar 5µm.

**Movie 4**

Live imaging of meiosis I in *hcp-2*-mutant worms expressing CLS-2::GFP::HCP-1^CTD^ fusion protein in indicated conditions. Microtubules (GFP::TBA-2^α-tubulin^) in green, DNA (mCherry::HIS-11^H2B^) in magenta. Time in seconds relative to anaphase I onset. Scale bar 5µm.

**Movie 5**

Live imaging of meiosis I in *hcp-2*-mutant worms expressing CLS-2^ΔCTD^::GFP::HCP-1^CTD^ fusion protein in indicated conditions. Microtubules (GFP::TBA-2^α-tubulin^) in green, DNA (mCherry::HIS-11^H2B^) in magenta. Time in seconds relative to anaphase I onset. Scale bar 5µm.

**Movie 6**

Live imaging of meiosis I in indicated transgenic worms upon depletion of endogenous *cls-2*. Microtubules (GFP::TBA-2^α-tubulin^) in green, DNA (mCherry::HIS-11^H2B^) in magenta. Time in seconds relative to anaphase I onset. Scale bar 5µm.

**Movie 7**

TIRF microscopy-mediated live imaging of in vitro microtubule (magenta) polymerization dynamics from GMPCPP seeds (cyan) in the presence of indicated protein (100 nm each). Scale bar 10 µm.

## SUPPLEMENTAL MATERIAL

**Supplemental Table S1**

List of *C. elegans* strains used in this study.

## SOURCE DATA

**Figure 1 – source data 1 - Figure 1B source data**

Source data for quantification of BUB-1 signal at kinetochores in *bub-3(ok3437)* mutants (*bub-3Δ*) and in control N2. Integrated intensity measurements corrected by integrated background noise (Integrated signal intensity/Average noise) of immunolocalized BUB-1 at kinetochores in fixed oocytes of *bub-3(ok3437)* mutants (*bub-3Δ*) compared to N2 wild type controls. Descriptive statistics and details of unpaired t-test for comparison of samples are provided.

**Figure 2 – source data 1 – Panel E source data**

Source data of embryonic viability assay of *hcp-2(ijm6)* mutant worms carrying *gfp::hcp-1* transgene variants, upon depletion of endogenous *hcp-1*.

**Figure 2 – Figure supplement 1 – source data 1 – Panel B source data**

Source data of embryonic viability assay of *hcp-2(ijm6)* mutant worms carrying *gfp::hcp-1* transgene variants, when endogenous *hcp-1* is not depleted.

**Figure 2 – Figure supplement 1 – source data 2 – Panel C source data**

Western blot raw images of GFP::HCP-1 fusion protein variants in worm extracts.

**Figure 3 – source data 1 – Panel A source data**

Quantification of cells showing meiotic defects and cells without meiotic defects in indicated conditions.

**Figure 3 – source data 2 – Panel B source data**

Quantification of spindle density and spindle size during meiosis I in indicated conditions. Raw quantifications, corrected intensities of GFP::TBA-2^α-tubulin^ signal (GFP Integrated Intensity/Background Noise), statistics and details of ANOVA multiple comparisons are provided.

**Figure 3 – Figure supplement 1 - source data 1**

Quantification of cells showing meiotic defects and cells without meiotic defects.

**Figure 4 – source data 1 – Panel B source data**

Quantification of cells showing meiotic defects and cells without meiotic defects.

**Figure 4 – source data 2 – Panel C source data**

Quantification of spindle density and spindle size during meiosis I in indicated *hcp-2(ijm6)* mutants upon depletion of endogenous *hcp-1*. Raw quantifications, corrected intensities of GFP::TBA-2^α-tubulin^ signal (GFP Integrated Intensity/Background Noise), statistics and details of Kruskal-Wallis multiple comparisons are provided.

**Figure 4 – source data 3 – Panel F source data**

Source data of embryonic viability assay of *hcp-2(ijm6)* mutant, *cls-2^FL^::gfp::hcp-1^CTD^* transgenic worms upon depletion of *hcp-1* and/or *cls-2*.

**Figure 4 – source data 4 – Panel G source data**

Quantification of cells showing meiotic defects and cells without meiotic defects in *hcp-2(ijm6)* mutant, *cls-2^FL^::gfp::hcp-1^CTD^*transgenic worms upon depletion of *hcp-1* and/or *cls-2*.

**Figure 4 – Figure supplement 1 – source data 1 – Panel A source data**

Quantification of cells showing meiotic defects and cells without meiotic defects in *bub-1^ΔKD^::mCherry* transgenic worms.

**Figure 4 – Figure supplement 1 – source data 2 – Panel E source data**

Quantification of cells showing meiotic defects and cells without meiotic defects in *hcp-2(ijm6)* mutant, *cls-2^ΔCTD^::gfp::hcp-1^CTD^*transgenic worms upon depletion of *cls-2* or both *hcp-*1 and *cls-2*.

**Figure 5 – source data 1 – Panel B source data**

Source data of embryonic lethality rescue assay of *cls-2::gfp* transgenic variants upon depletion of *endogenous cls-2*.

**Figure 5 – source data 2 – Panel D source data**

Quantification of cells showing meiotic defects and cells without meiotic defects in *cls-2::gfp* transgenic variants upon depletion of *cls-2*.

**Figure 5 – Figure supplement 1 – source data 1 - Panel D source data**

Western blot raw images of GFP::CLS-2 fusion protein variants in worm extracts.

**Figure 6 – Figure supplement 1 – source data 1 – Panel D source data**

Source data of embryonic viability assay of worms *carrying a cls-2::gfp* transgene with mutations in the SR-rich domain, upon depletion of *cls-2* compared to non-depleted controls.

**Figure 6 – source data 1 – Panel C-F source data**

Source data for quantification of microtubule dynamics parameters. Parameters were extracted from kymographs using the ImageJ software (https://imagej.nih.gov/ij/index.html). Each sheet corresponds to the measure of a single microtubule dynamics parameter: growth, shrinkage rates and durations, and pause number and durations extracted from kymographs. The first sheet (read me) provide details about data extraction.

**Figure 6 – source data 2 – Panel D-F source data**

Measure (counting) of catastrophe, rescue and pause events from the kymographs analyzed in Figure 6.

## DATA AVAILABILITY

All data generated or analysed during this study are included in the Source Data file.

---

**Figure 2 - Figure supplement 1.**
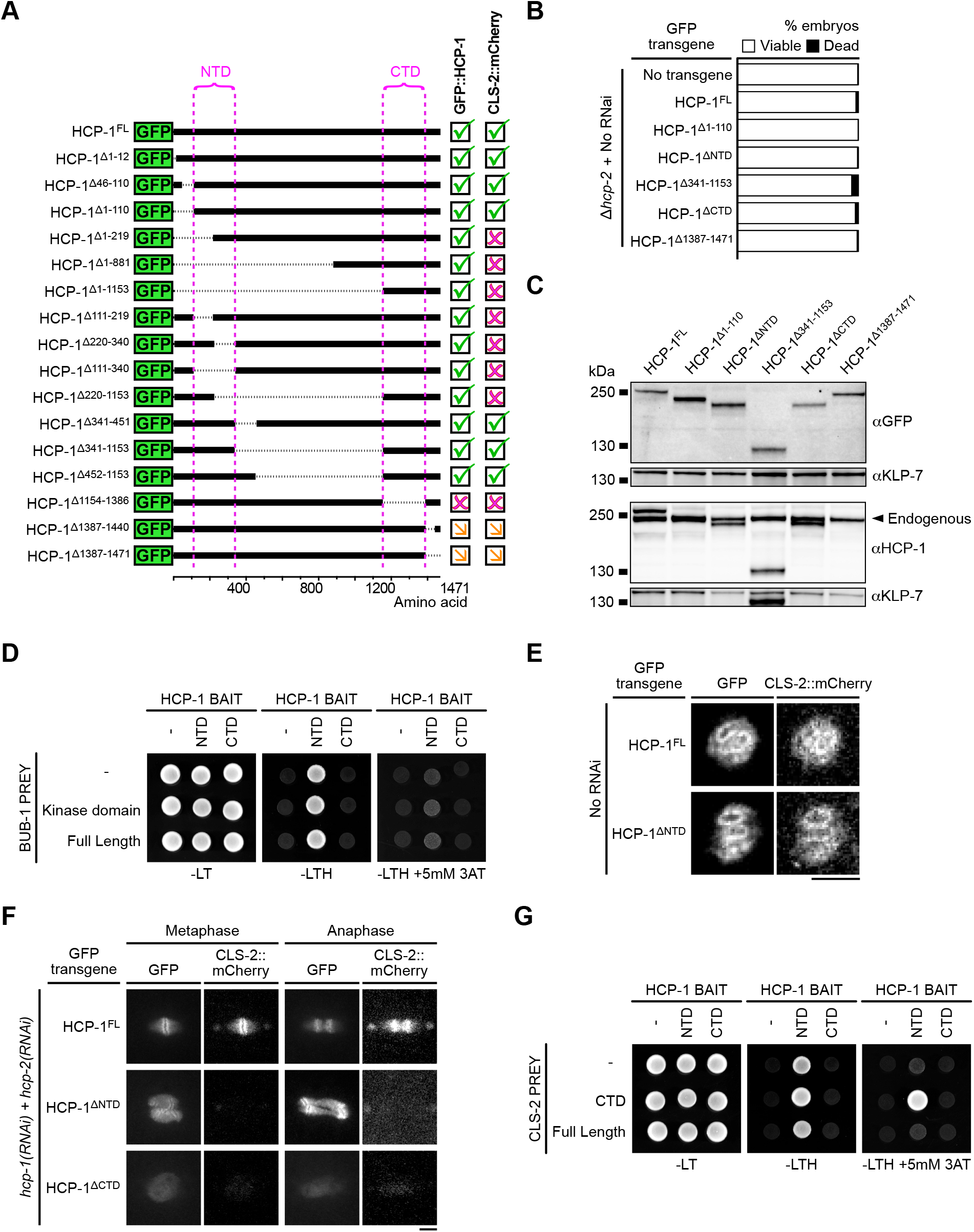
(A) Schematics of truncated GFP::HCP-1 fusions used to identify the NTD and CTD regions. Indicated on the right, presence (tick), absence (cross) or decrease (arrow) of GFP::HCP-1 and CLS-2::mCherry signals in oocytes and zygotes. (B) Embryonic viability assay of *hcp-2(ĳm6) (Δhcp-2)* mutants in indicated conditions. (C) Western blot of full-protein extracts (50 worms per lane). Arrowhead shows endogenous HCP-1. (D) Yeast two hybrid interaction assay between HCP-1 domains (baits) and CLS-2 (preys). (E,F) Localization of CLS-2::mCherry in meiosis I (E) and during the first mitotic division of the zygote (F) in indicated conditions (n≥10). Scale bars, 5µm. (F) Yeast two hybrid interaction assay between HCP-1 domains (baits) and BUB-1 (preys).

**Figure 3 - supplement 1.**
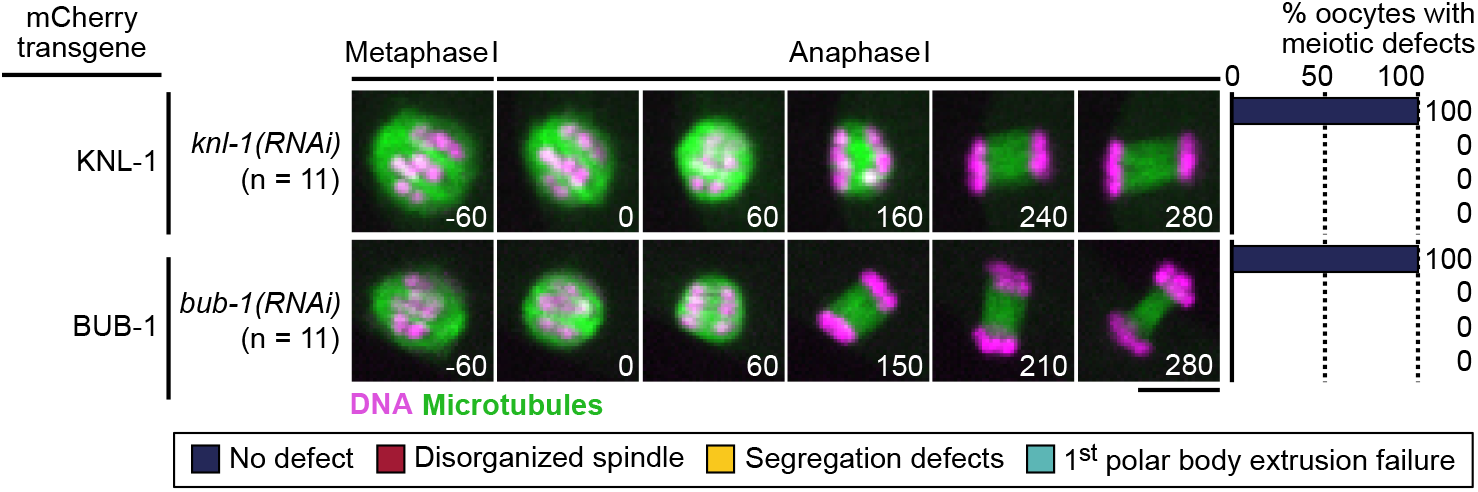
Stills from live imaging of *knl-1::mCherry*, *bub-1::mCherry* and transgenic oocytes in indicated conditions. Microtubules (GFP::TBB-2^ß-tubulin^) in green, chromosomes (mCherry::HIS-11^H2B^) in magenta. Right panels indicate quantifications of meiotic defects shown in the color key. Time in seconds relative to anaphase I onset. Scale bar, 5 µm.

**Figure 4 – Figure supplement 1.**
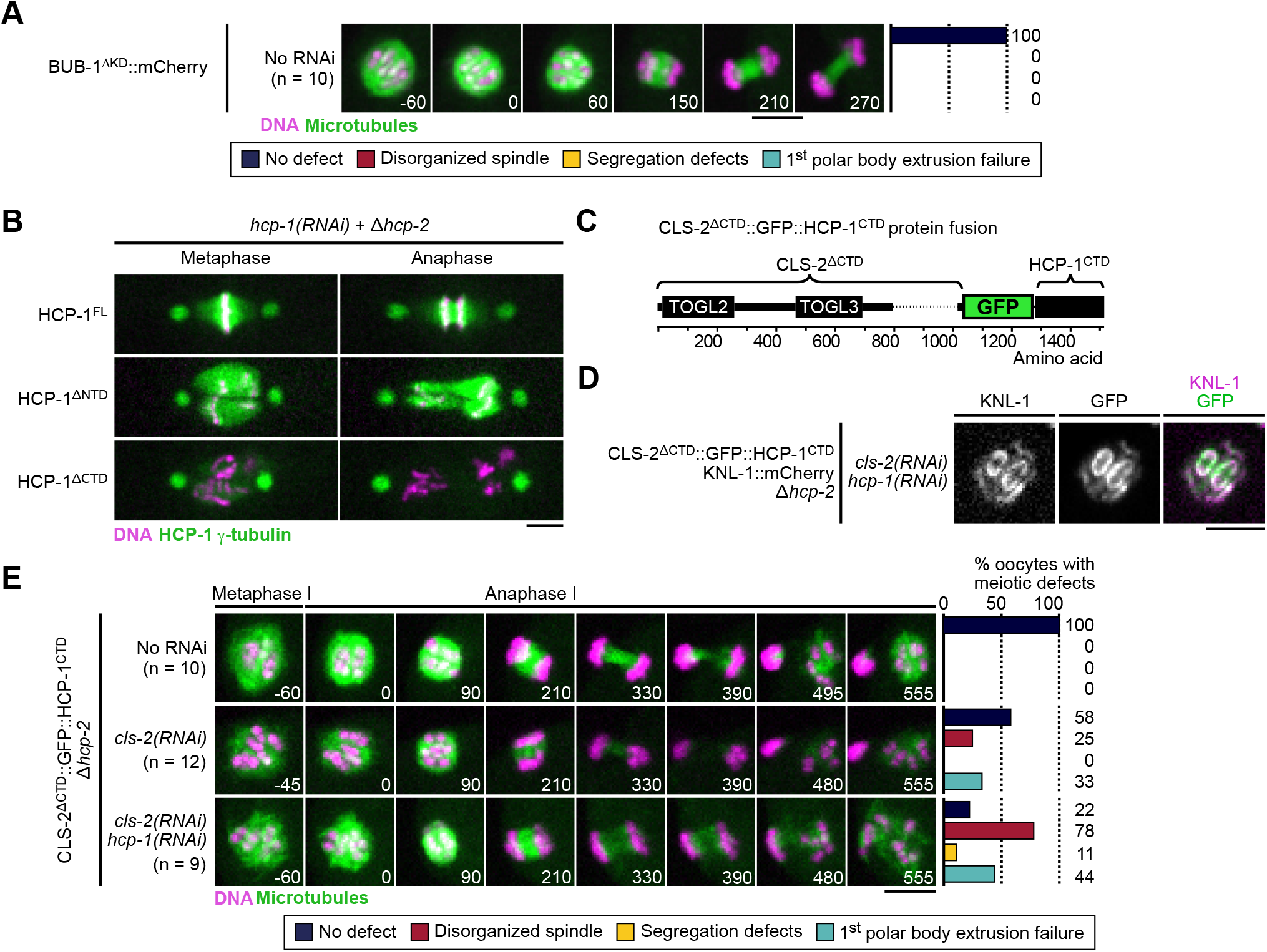
(A) Stills from live imaging of *bub-1ΔKD::mCherry* transgenic oocytes without silencing of endogenous *bub-1*. Microtubules (GFP::TBB-2^ß-tubulin^) in green, chromosomes (mCherry::HIS-11^H2B^) in magenta. Right panels indicate quantifications of meiotic defects shown in the color key. Time in seconds relative to anaphase I onset. (B) Localization of indicated GFP::HCP-1 fusion proteins (green) during the first zygotic mitosis (n≥9). Centrosomes (GFP::TBG-1^ɣ-tubulin^) in green, chromosomes (mCherry::HIS-11^H2B^) in magenta. (C) Schematics of the CLS-2^ΔCTD^::GFP::HCP-1^CTD^ protein fusion. (D) Localization of CLS-2^ΔCTD^::GFP::HCP-1^CTD^ in metaphase I, relative to KNL-1 marked kinetochores (KNL-1::mCherry, magenta), in *Δhcp-2* mutants depleted of endogenous *hcp-1* and *cls-2* (n=11). (E) Stills from live imaging of meiosis I (left) and color-coded meiotic defects scored (right) in CLS-2^ΔCTD^::GFP::HCP-1^CTD^, *Δhcp-2* worms in indicated conditions. Time in seconds relative to anaphase I onset. Scale bars, 5 µm.

**Figure 5 – Figure supplement 1.**
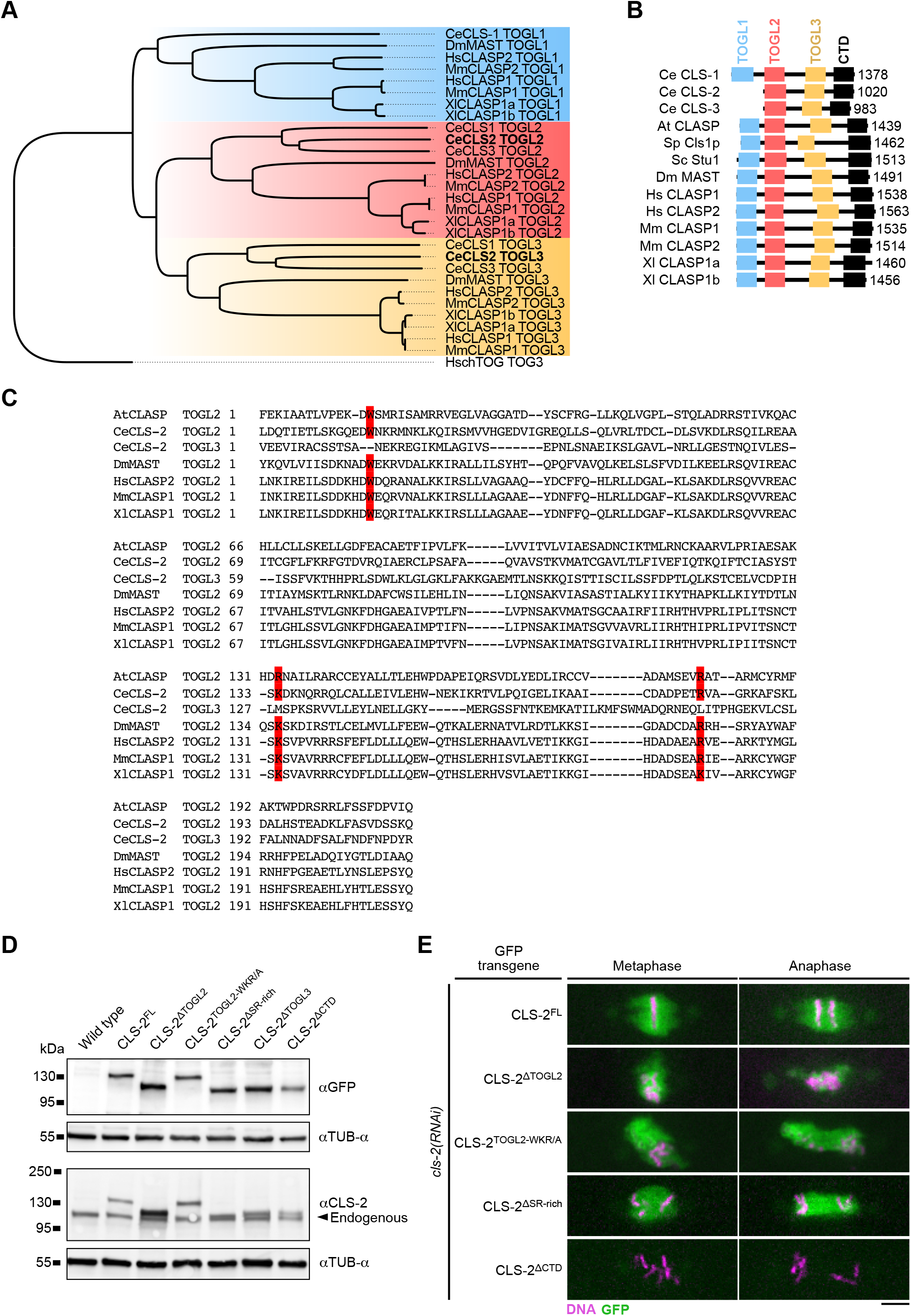
(A) Maximum Likelihood phylogenetic tree of eukaryotic CLASP TOG-like domains, rooted on human chTOG TOG3. (B) Comparative schematics of eukaryotic CLASP protein structures. Sizes in amino-acids are indicated on the right. (C) Protein sequence alignment of indicated eukaryotic TOGL domains. Conserved amino acids mutated in the CLS-2^TOGL2-WKR/A^ mutant in red. (D) Western blot of full-worm protein extracts (50 worms per lane). Arrowhead shows endogenous CLS-2. (E) Mitotic localization of indicated CLS-2::GFP fusion proteins upon depletion of endogenous *cls-2* (green). DNA (mCherry::HIS-11) in magenta (n≥10). Scale bar, 5 µm.

**Figure 6 – Figure supplement 1.**
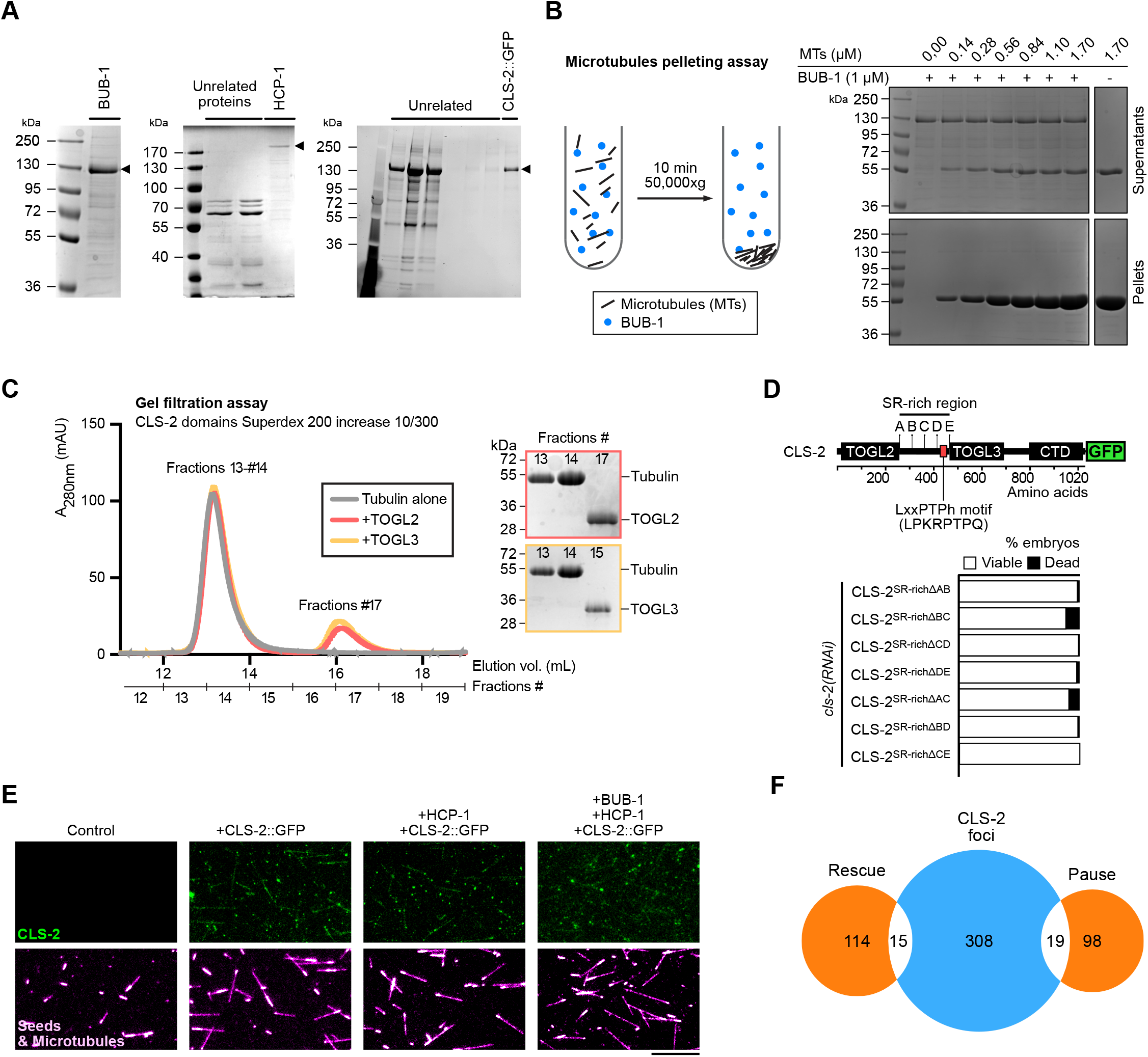
(A) Coomassie-stained gels of purified proteins used for *in vitro* assays. Arrowheads indicate the protein of interest. (B) Microtubule pelleting assay in the presence of 1 µM purified BUB-1 protein. Schematics of the experiment principle (left) and Coomassie staining of supernatant and pellet fractions (right) with indicated concentrations of microtubules (MTs). (C) Gel filtration assay of tubulin in the presence of indicated CLS-2 TOGL domains. Coomassie staining of the fractions of interest are shown on the right. (D) Embryonic viability assay of worms carrying *cls-2::gfp* transgenes with indicated truncations in the SR-rich region of CLS-2 upon depletion of endogenous *cls-2.* Position of the divergent LxxPTPh motif is indicated on the protein fusion diagram. (E-F) TIRF-microscopy localization of 100 nM purified CLS-2::GFP protein (green) on microtubules (magenta). Stills from imaging in indicated conditions (E) and Venn diagram of CLS-2::GFP foci associated with microtubule rescue and pause events (F). Scale bar 10 µm.

